# Unveiling the cell biology of hippocampal neurons with dendritic axon origin

**DOI:** 10.1101/2023.09.14.557747

**Authors:** Yuhao Han, Roland Thuenauer, Kay Grünewald, Marina Mikhaylova

**Affiliations:** AG Optobiology, Institute of Biology, Humboldt Universität zu Berlin, Berlin, 10115 Germany; Guest Group “Neuronal Protein Transport”, Centre for Molecular Neurobiology, University Medical Center Hamburg-Eppendorf, Hamburg, 20251 Germany; Centre for Structural Systems Biology, Hamburg, 22607 Germany; Structural Cell Biology of Viruses, Leibniz Institute of Virology (LIV), Hamburg, 20251 Germany; Advanced Light and Fluorescence Microscopy (ALFM) Facility, Centre for Structural Systems Biology, Hamburg, 22607 Germany; Technology Platform Light Microscopy, University of Hamburg, Hamburg, Germany; Department of Chemistry, University of Hamburg, Hamburg, 20146 Germany; Technology Platform Microscopy and Image Analysis (TP MIA), Leibniz Institute of Virology (LIV), Hamburg, Germany

**Keywords:** Axon-carrying dendrite neurons, axon initial segment (AIS), cytoskeleton, AIS plasticity, cisternal organelle

## Abstract

In mammalian axon-carrying-dendrite (AcD) neurons, the axon emanates from a basal dendrite, instead of the soma, to create a privileged route for action potential generation at the axon initial segment (AIS). However, it is unclear how such unusual morphology is established and whether the structure and function of the AIS in AcD neurons is preserved. Here, we show that the AcD neurons follow an intrinsically encoded developmental program where a single precursor neurite first gives rise to the axon and then to the AcD. The AIS possesses a similar cytoskeletal architecture as the canonical AIS that stems from the soma, and similarly functions as a trafficking barrier to retain axon-specific molecular composition. However, unlike soma-derived AIS, the AIS of AcD neurons does not undergo homeostatic-plasticity, contains less cisternal organelles and receives fewer inhibitory inputs. These distinct features of the AIS could account for the higher intrinsic excitability of AcD neurons.

**Highlights:**

- The development of AcD neurons is intrinsically driven and does not depend on specific connectivity patterns
- AcD neurons generate the axon and AcD from the same precursor neurite
- The stem dendrite of an AcD neuron displays axon-like microtubule organization
- Similar to the AIS emerging from the soma, the AIS of AcD neurons can selectively filter dendritic cargo
- The AIS of AcD neurons does not undergo chronic homeostatic plasticity and receives fewer inhibitory inputs

## Introduction

The highly polarised morphology of neurons enables the directional transmission of electrical signals, a fundamental mode of communication within neuronal circuits. Typically, mammalian brain neurons have a single axon and multiple highly branched dendrites that emerge directly from the soma. Synaptic inputs are received at dendrites and propagate along them towards the soma in the form of membrane depolarisation. Following somatic integration and upon reaching a specific threshold at the axon initial segment (AIS), these inputs trigger the generation of action potentials (APs) – a crucial mechanism enabling neurotransmission.

In addition to such a classical view of neuronal morphology, it has been reported that the axon of a neuron may not necessary originate from the soma but can also emerge from a basal dendrite. Already in 1899, Ramon Cajal observed such neurons in invertebrate abdominal ganglia.^1^ Later, neurons with dendritic axon origin were also discovered in the neocortex and hippocampus of mammalian brains, including human.^2–4^ Given their axon origin, this type of neurons is referred to as axon-carrying dendrite (AcD) neurons. A typical feature of AcD neurons is that the axon and the AcD branch out from a common dendritic root, called stem dendrite.^2–4^ In the rodent hippocampus, AcD neurons were found to be primarily distributed at the central part of the CA1 region, where they represent over ∼50% of the entire excitatory neuron population.^3^ Electrophysiological experiments suggested that, in AcD neurons, inputs arriving at the AcD would have a higher probability to trigger APs than the regular somatic dendrites.^3^ Since the axon is adjacent to the AcD, synaptic inputs from the AcD can bypass somatic integration, and directly flow into the axon as APs.^3^ This novel AcD-to-axon paradigm of AP transmission was shown to allow AcD neurons evade global peri-somatic inhibition, and therefore being selectively activated during sharp-wave ripples, a process associated with memory consolidation.^4^ Collectively, these findings revealed that the dendritic axon origin has a substantial impact on the electrophysiological behaviour of hippocampal excitatory neurons. Despite their functional importance, the cellular features of AcD neurons and the developmental sequence leading to such morphology have remained unexplored.

The central dogma of neuron development is that neurons break the symmetry by initially forming the axon. After mitosis, the spherical-shaped neurons first generate a pool of nascent neurites and then specify a single precursor neurite to become an axon.^5,6^ Once the axon is properly established, the remaining neurites begin to differentiate into dendrites and synaptic connections between neurons are eventually formed.^5,6^ Such development program is mostly genetically encoded and can be reproduced in dissociated cell cultures.^5^ However, the timing, topology of subcellular domains, connectivity, and certain other specific aspects of neuron development can vary between cell types. This variation may be attributed to the distinct microenvironments formed by *in vivo* guidance molecules.^6,7^ The morphology of AcD neurons poses a great challenge to this canonical sequence of neuron development. To date it is unclear whether AcD neurons adhere to classical developmental stages. It is yet to be determined whether transformation into the AcD type is intrinsically driven or if it depends on specific connectivity patterns and extracellular factors. Answering these questions will provide great insights into the diversity of neuronal development programs.

The cytoskeleton plays an instrumental role in the establishment and maintenance of neuronal axo-dendritic polarity. During development, the cytoskeleton of the pre-mature axon acquires distinctive characteristics, including the uniform plus-end-out orientation of microtubules (MTs). Axonal MTs are also stabilised by accumulating specific post-translational modifications (PTMs) of tubulins, such as acetylation.^8–10^ Contrarily, dendrites are characterized by a lower presence of acetylated microtubules (MTs) compared to axons, and a mixed orientation of MT plus ends.^8,10,14^ The unique axonal MT organisation together with PTMs facilitates the growth and maturation of the axon by enabling targeted delivery of vesicles containing axon-specific proteins and lipids via kinesin and dynein motor proteins.^11–13^ It is an open question how the targeting of axonal cargo could work in AcD neurons when axonal vesicles first have to pass dendritic compartment before entering the axon.

The AIS is another important axonal domain which is present in the proximal part of the axon and extending about 20-60 µm towards the distal end. The molecular organisation of the AIS is well-tailored for its function and specialized cytoskeletal components can only be found in this compartment. Thus, individual MTs within the AIS are interconnected by the AIS-specific microtubule-associated protein (MAP) TRIM46 to form MT fascicles.^15^ The AIS-specific scaffolding proteins ankyrin-G (AnkG) and βIV-spectrin are localised in between periodically organised rings of filamentous actin (F-actin) that define the so-called membrane-associated periodic cytoskeleton (MPS).^16–19^ F-actin additionally forms patches.^20–22^ Together, such specialised cytoskeleton provide a scaffolding platform for: 1) anchoring membrane proteins, such as cell adhesion molecules (CAMs) and voltage-gated ion channels (VGICs)^16,19^; 2) enables the formation of AIS-specific endoplasmic reticulum (ER) specialisations called cisternal organelles (COs), which is mediated by the actin binding protein synaptopodin (synpo) and is responsible for Ca^2+^ handling^23–25;^ and 3) allows for the formation of an AIS-specific extracellular matrix (ECM)^19,26^. The AIS serves not only as a molecular barrier to exclude somatodendritic proteins^20–22^, but also to initiate action potentials along the axon, thereby regulating neuronal excitability and homeostasis.^16,19,27,28^ It has been shown that the AIS of excitatory neurons undergo structural remodelling to compensate for high neuronal activity, a process known as AIS plasticity.^27,28^ Furthermore, the AIS of excitatory neurons is innervated by inhibitory synapses for fine-tuning of excitability during network activities^29^. How similar the AIS features of AcD neurons are to neurons with a somatic axon origin is currently unknown.

In this study, we seek to provide a comprehensive cell biological profile of hippocampal AcD neurons by focusing on the developmental processes of these neurons and providing a detailed characterization of the axon initial segment (AIS), including its structural and functional properties, under both basal and enhanced neuronal activity conditions.

## Results

### Development of AcD neurons does not require specific neuronal connectivity patterns and extracellular guidance

To investigate the development of AcD neurons, we used dissociated hippocampal primary neurons as model system. An advantage of such system is that it lacks the *in vivo* microenvironments that essentially guide neuron development. We hypothesized that if dissociated neurons have an axon emanating from the AcD, their development must be driven by genetically encoded factors rather than specific connectivity patterns or a gradient of guidance molecules that instruct neuronal migration and brain development. Interestingly, immunofluorescent staining of primary neurons with the AIS marker AnkG and the somatodendritic marker MAP2 clearly showed that a subset of cells display the AcD morphology, as the AIS is branched out from a dendrite and located further away from the soma (**Figure 1A**). Already at 3 days *in vitro* (DIV), AcD neurons were present and the AnkG fluorescent signal began to accumulate at the proximal axon region (**Figure 1A**). AcD neurons were also found in cultures fixed at later time point (DIV5, 7 and 12) (**Figure 1A**), and the AnkG signal appeared to be more continuous and evident, as the AIS become more mature (**Figure 1A**). The same time line of AIS formation was also observed in nonAcD neurons (**Figure 1A**), which is consistent with a previous study.^30^ Together, these results suggest that the development of AcD neurons is indeed an intrinsically driven process, and that the dendritic axon origin has no impact on the timing of AIS assembly.

**Figure 1.**
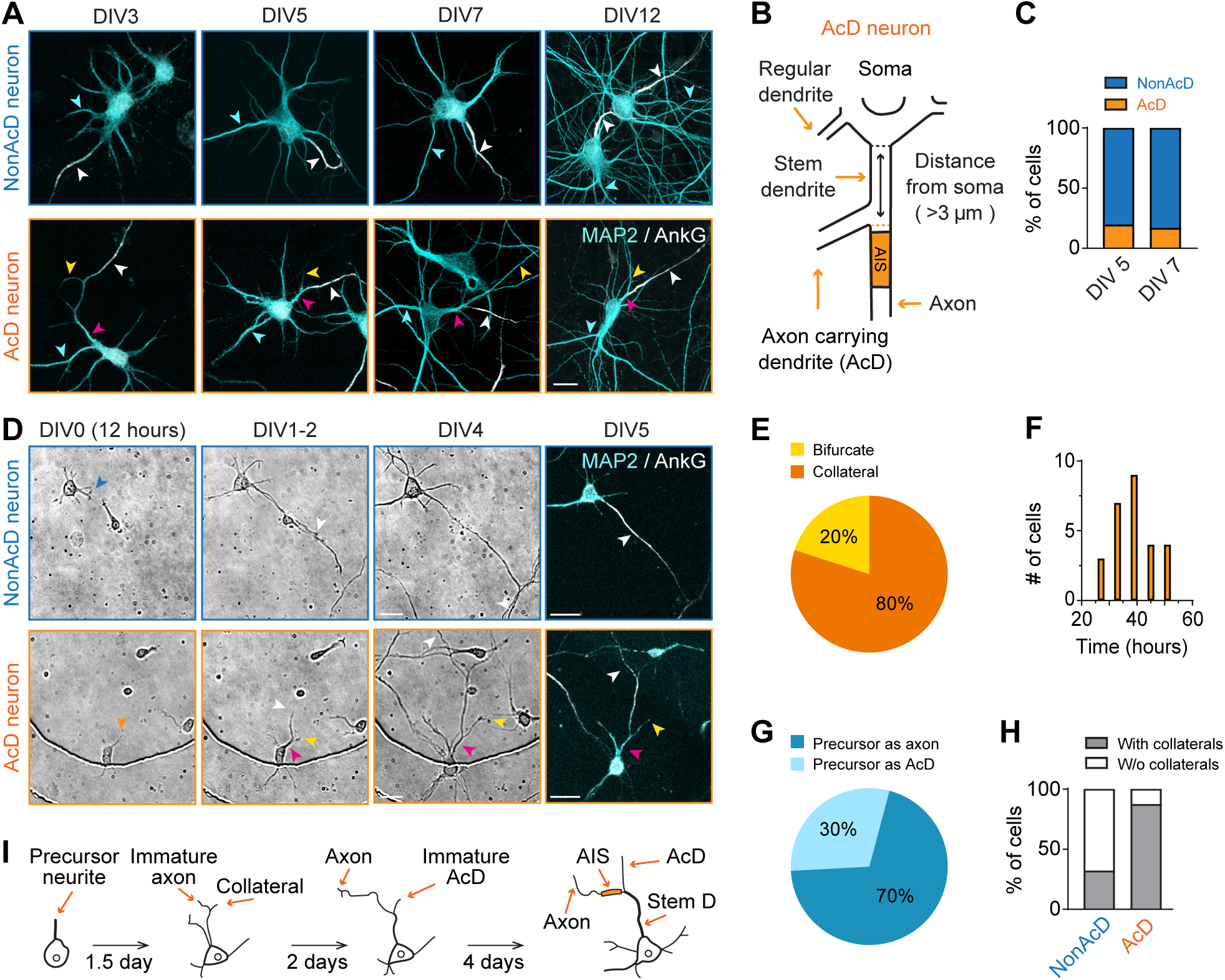
Development of neurons with axon carrying dendrite. **(A)** Representative images of dissociated hippocampal primary neurons with nonAcD (upper row) and AcD (lower row) morphology at different time points. AnkG and MAP2 immunostaining indicate the axon/AIS and dendrites of a neuron, respectively. White arrowhead indicates AIS; yellow and pink arrowhead indicates AcD and stem dendrite of AcD neuron respectively; cyan arrowhead indicates regular dendrite. Scale bar is 20 µm. **(B)** Schematic of AcD neuron. Black dashed line indicates the edge of soma; orange dashed line indicates the beginning of an axon. Black solid line with double arrowheads indicate distance from the edge of soma to beginning of axon. **(C)** Percentage of nonAcD and AcD neurons in dissociated culture at DIV5 and DIV7. n = 6 coverslips from 3 independent cultures, DIV5: 83 AcD neurons out of 427 cells, DIV7: 47 AcD neurons out of 387 cells. **(D)** Time-lapse images of nonAcD (upper row) and AcD neurons (lower row) at different developmental stages. The precursor neurite in the displayed AcD neuron became an axon and collateral from the precursor neurite developed as AcD. Orange arrowhead indicates the precursor neurite of AcD neuron; blue arrowhead indicates precursor neurite of nonAcD neuron; white arrowhead indicates axon; yellow and pink arrowhead indicates AcD and stem dendrite of AcD neuron, respectively. Scale bar is 20 µm. Cell body with dendrites and the AIS are labelled by MAP2 and AnkG staining, respectively. Related to **Video S1** and **S2**. **(E)** Percentage of AcD neurons generate collateral at or bifurcate. 1 culture preparation, nonAcD: n= 25 cells, AcD: n= 27 cells. **(F)** Time points of a collateral formation. 1 neuronal culture preparation, nonAcD: n= 25 cells, AcD: n= 27 cells. **(G)** Percentage of AcD neurons where a precursor neurite developed into the axon or AcD. 1 culture preparation, AcD: n= 23 cells. **(H)** Percentage of nonAcD and AcD neurons that formed collaterals at the proximal region of the precursor neurite during development. 1 culture preparation, nonAcD: n= 25 cells, AcD: n= 23 cells. **(I)** Proposed model of AcD neuron development.

For further investigation, we classified hippocampal neurons into AcD and nonAcD categories based on previously published standards^2–4^ (**Figure 1B**; **Figure S1D**; see methods for details). We considered neurons with an axon emerging from a MAP2-positive dendrite at the distance longer than 3 µm away from the soma as AcD neurons. Otherwise, neurons were considered as nonAcD. We quantified the percentage of AcD neurons in dissociated cultures at DIV5 and DIV7, at which neuronal polarity has been established. Quantification showed that AcD neurons make up 15-20% of the entire population at both time points (**Figure 1C**), indicating that dissociated cultures provide a suitable model for investigating the cell biology of AcD neurons.

### AcD neurons follow consensus developmental sequence and mostly generate the AcD as axonal collateral

The canonical process of neuron development *in vitro* is defined into five stages.^5^ After surface attachment, neurons begin to generate a pool of nascent neurites and then specify a single precursor neurite as the axon (stage 1-3; DIV0-3; **Figure 1D**; **Video S1**). After axon formation, the remaining neurites begin to differentiate into the dendrites (stage 4; DIV4-7; **Figure 1D**; **Video S1**), followed by the establishment of synaptic contacts (stage 5; after DIV7). Here we asked whether AcD neurons follow the same developmental sequence, given their atypical axon origin. To test this, we conducted continuous time-lapse microscopy recordings of dissociated neurons from DIV0 to DIV5 and subsequently performed immunostaining with AnkG and MAP2 for post-hoc categorization of neuronal types. Time-lapse imaging data revealed that AcD neurons indeed also first grow an axon from a precursor neurite and then establish the dendrites (**Figure 1D**; **Figure S1A**; **Video S2**). Notably, during early developmental stages (DIV1-2), a collateral neurite was generated from the proximal region of the precursor neurite that is designated as the axon (**Figure 1D**; **Figure S1A**; **Video S2**). This collateral underwent several rounds of elongation and retraction while the precursor neurite kept developing as the axon (**Figure S1A**; **Video S2**). Eventually, once the axon was defined, the collateral started to mature as the AcD and the former proximal region of the axon precursor neurite was transformed into the stem dendrite (**Figure 1D**; **Figure S1A**; **Video S2**).

Quantitative analysis showed that about 80% of the AcD neurons generated collateral at the proximal region of the precursor neurite (**Figure 1E**), and the average time point of collateral formation was at 39 hours (DIV1.5) after plating (**Figure 1F**). About 20% of the AcD neurons bifurcated the precursor neurite growth cone to form the axon and AcD (**Figure 1E**; **Figure S1B**; **Video S3**). Among the 80% of AcD neurons that formed collateral at the precursor neurite, the majority of the population (70%) designated the precursor neurite as the axon and formed the collateral as AcD (**Figure 1G**). Only a smaller population (30%) of neurons instead developed the collateral as the axon, and eventually turned the precursor neurite into the AcD (**Figure 1G**; **Figure S1A**; **Video S4**). We also assessed the collateral genesis at the proximal region of the axon precursor neurite in nonAcD neurons. We found that the percentage of nonAcD neurons that formed collaterals at DIV1 is much lower than for AcD neurons (30% nonAcD neurons vs. 80% AcD neurons; **Figure 1H**), and these branches completely retracted at DIV4 (**Figure S1C**). Collectively, our findings suggest that AcD neurons still follow consensus developmental steps like nonAcD neurons to first establish the axon and then the dendrites (**Figure 1I**). AcDs are derived mainly from a branch formed at the basal region of immature axons during early development (**Figure 1I**). Although, multiple alternative strategies do exist during AcD neuron development which is similar to what has been shown for nonAcD neurons in vitro.^5^

### The stem dendrite of AcD neurons has axon-like microtubule organisation

The differences in MT orientation and post-translational modifications between axons and dendrites play a crucial role in guiding selective cargo transport in neurons.^8,9,31,32^ We next asked whether AcD neurons have similar MT orientations and types of MT PTMs in the neurites as nonAcD neurons. We were particularly interested in the stem dendrite, as this region connects the soma and axon. To track MT plus ends, we moderately overexpressed MT plus end binding protein-3 tagged with Tdtomato (EB3-Tdtomato) in mature (DIV14) dissociated hippocampal cultures. For axon identification, we live-labelled the AIS with a fluorescently conjugated antibody against the extracellular domain of the AIS enriched membrane protein neurofascin (anti-NF-CF640R) (**Figure S1E**). We found that the AIS, axon and regular somatic dendrites of AcD neurons displayed MT orientations similar to nonAcD neurons. As indicated by EB3-Tdtomato trajectories, in both AcD and nonAcD neurons, MT plus ends in the AIS and axon are uniformly oriented towards the distal part of the axon but are anti-parallel in the regular somatic dendrites (**Figure 2A** and **2C**; **Video S5 - S7**). We also observed the typical dendritic MT orientation in the AcD, where the MT plus ends grew towards both directions (**Figure 2B** and **2C**; **Video S5**). By stark contrast, we found that in the stem dendrite of AcD neurons, nearly 90% of the EB3-Tdtomato trajectories showed a unidirectional movement either towards the axon or the AcD (**Figure 2B** and **2C**; **Video S5**). This unidirectional plus end-out MT orientation in the stem dendrite highly resembles that of axons.

**Figure 2.**
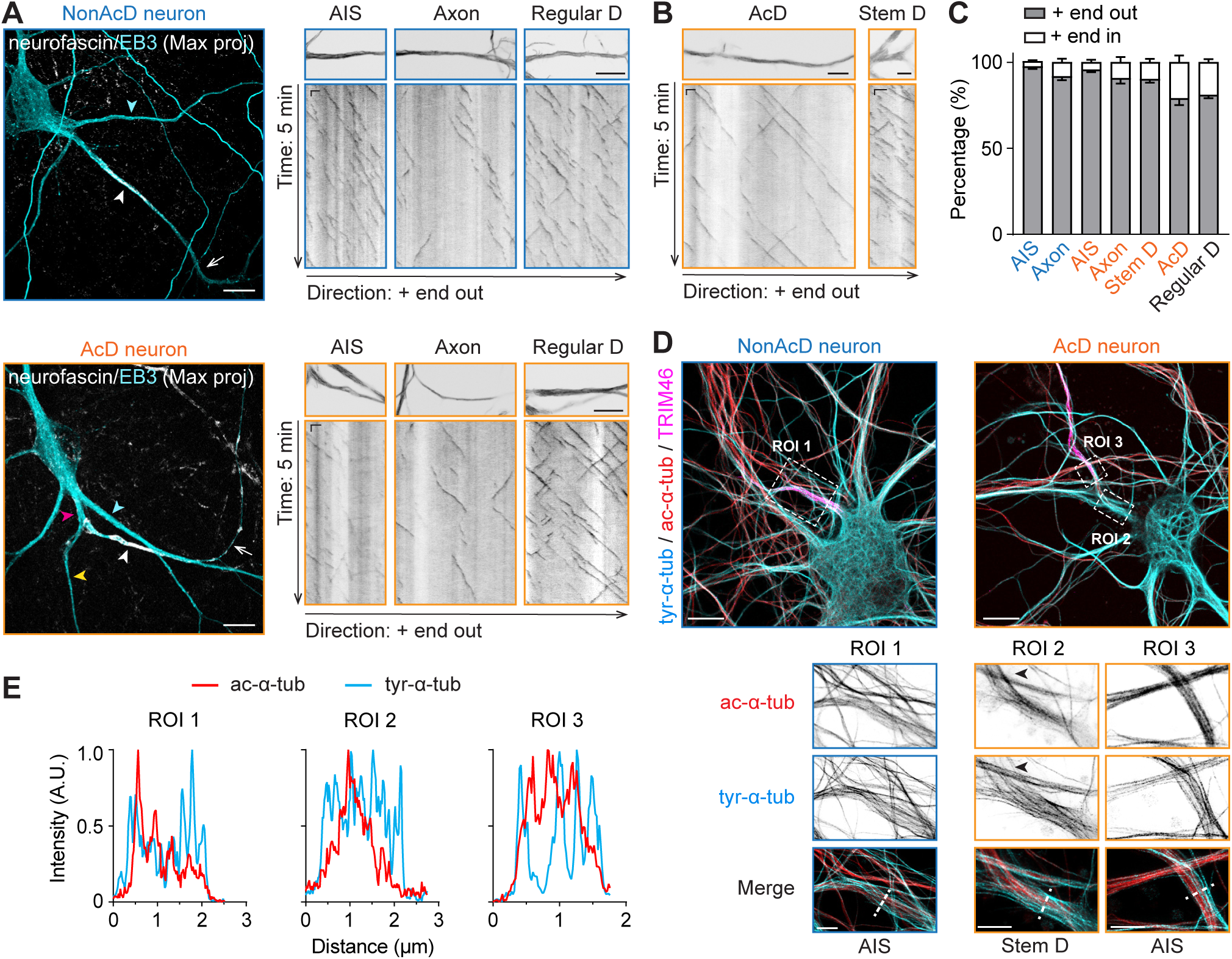
The stem dendrite of AcD neuron exhibits axon-like MT organisation. **(A)** Left panel: Maximum intensity projection of DIV14 nonAcD and AcD neurons transfected with EB3-Tdtomato for visualisation of MT plus ends. The AIS is live-labelled with anti-NF-CF640R antibody. White arrowhead indicates the AIS and white arrow indicates the axon. Cyan arrowhead indicates regular dendrite (Regular D); yellow and pink arrowhead indicates AcD and stem dendrite of AcD neuron, respectively. Scale bar is 10 µm. Right panel: 5 minutes time projection and kymograph of EB3-Tdtomato in the AIS, axon and regular dendrite (Regular D) of nonAcD and AcD neuron shown in the left panel. Scale bar is 10 µm. Kymograph scale is 10 s (vertical) and 2 µm (horizontal). Related to **Video S5**, **S6** and **S7**. **(B)** 5 minutes time projection and kymograph of EB3-Tdtomato in AcD and stem dendrite (Stem D) of AcD neuron shown in (A). Scale bar is 5 µm. Kymograph scale is 10 s (vertical) and 2 µm (horizontal). Related to **Video S5**. **(C)** Percentage of MT plus end orientations in different regions of nonAcD and AcD neurons. Mean ± SEM, 3 independent cultures, AIS (nonAcD) n = 23, AIS (AcD) n = 17, axon (nonAcD) n = 21, axon (AcD) n= 15, stem dendrite (Stem D) n = 19, AcD n = 17, regular dendrite (Regular D) n = 67. **(D)** Top row: Representative confocal images of DIV10 nonAcD and AcD neurons stained for tyrosinated and acetylated tubulin, and the AIS marker TRIM46. Scale bar is 10 µm. Bottom row: Single plane 2D gSTED image of tyrosinated and acetylated MTs corresponding to ROIs in top row. Scale bar is 2.5 µm. ROI1 is the axon (AIS region) of the displayed nonAcD neuron. ROI2 and ROI3 is the stem dendrite and axon (AIS region) of the displayed AcD neuron, respectively. Black arrowhead in ROI2 indicates the axon. **(E)** Intensity profile of tyrosinated and acetylated MTs indicated by white dashed lines in ROIs shown in (**D**) bottom row.

Next, we performed gated-stimulation emission depletion (gSTED) super-resolution microscopy to visualise MT PTMs in the axon and stem dendrite of AcD neurons. Specifically, we looked into MT tyrosination and acetylation, which are the most common PTMs that represent dynamic and stable MTs, respectively. Previous studies suggested that acetylated and tyrosinated MTs have different spatial arrangements in the neurite^10,33^. Furthermore, the acetylated MTs are more enriched in neuronal axon^8^, and this enrichment of acetylated MTs instruct transport of axonal proteins by preferentially enabling vesicles driven by the kinesin-1 motor protein.^12,32^ Our data showed that both acetylated and tyrosinated MTs are present in the axon of AcD neurons (**Figure 2D**). The acetylated MTs are placed near the central core of the axon and surrounded by tyrosinated MTs (**Figure 2D** and **2E**). This spatial arrangement of MTs is the same as in the axon of nonAcD neurons (**Figure 2D** and **2E**) and is also consistent with the previous study^10^. Of note, we observed that acetylated MTs in AcD neurons were already present in the stem dendrite. Similarly to the axon, they were enveloped by tyrosinated MTs (**Figure 2D** and **2E**). Moreover, bundles of acetylated MT seem to extend from the stem dendrite directly into the axon, while they occur less in the AcD (**Figure 2D**). Altogether, our data suggest that MTs in AcD neurons are similarly organised as in nonAcD neurons, but the stem dendrite of AcD neurons likely inherits axonal MT organisation from its development to achieve targeted transport of axonal proteins.

### The AIS of AcD neurons has similar cytoskeletal organisation as nonAcD neurons

The AIS contains distinct cytoskeletal structures that help neurons to maintain polarity by segregating dendritic and axonal proteins^16,19^. We therefore proceeded with characterising the cytoskeletal structure of AcD neuron’s AIS. An important feature of the AIS MTs is their organisation into fascicles mediated by TRIM46. Similarly to nonAcD neurons, we found that in AcD neurons TRIM46 signal was clearly present and accumulated in the AIS (**Figure 3A**; **Figure 2D**). Another distinctive cytoskeletal feature of the AIS is the organisation of F-actin which is arranged into the MPS^16,34^, and F-actin patches^20–22^. To visualise F-actin at the AIS, we labelled dissociated neurons with phalloidin and AnkG, and performed gSTED imaging. We found that F-actin at the AIS of AcD neurons also formed patches and periodically arranged ring-like structures (**Figure 3B**). Similarly to nonAcD neurons, the distance between F-actin rings in AcD neuron’s AIS was ∼190 nm (**Figure 3C** and **3D**), suggesting there is no substantial difference in the actin cytoskeleton of AcD neuron’s AIS. Together, our data indicate that the actin and MT cytoskeleton in AcD neuron’s AIS is similarly organised as in nonAcD neurons.

**Figure 3.**
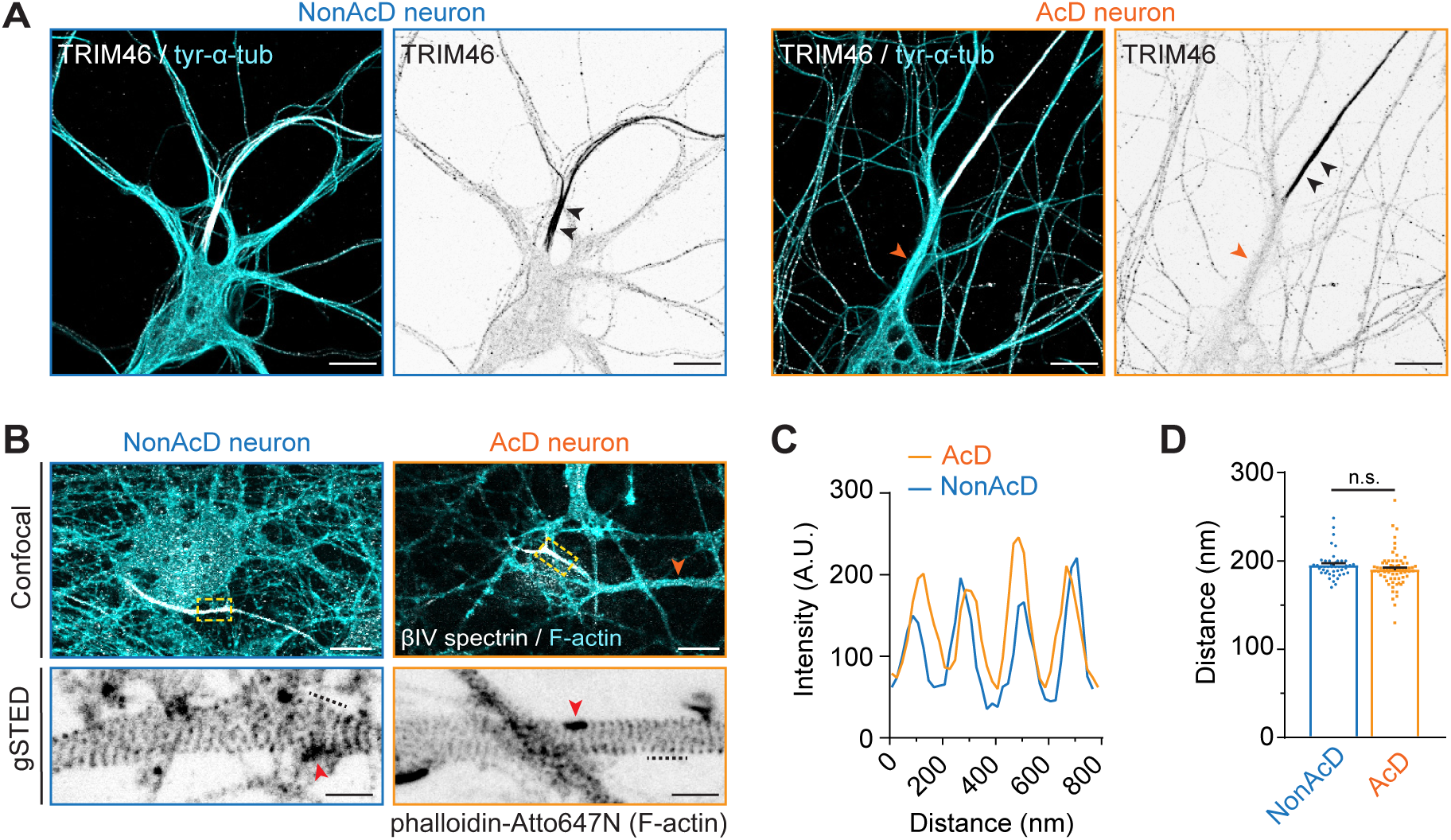
AcD neurons have similar MT and F-actin organisation at the AIS as in nonAcD neurons. **(A)** Representative images of AIS-specific MT bundles mediated by TRIM46 at the AIS of DIV10 nonAcD and AcD neurons. Orange arrowhead indicates the stem dendrite of the displayed AcD neuron. Black arrowheads indicate the AIS. Scale bar is 10 µm. **(B)** Top row: Representative confocal images of DIV14 nonAcD and AcD neurons stained with the AIS marker βIV-spectrin and F-actin probe Phalloidin-Atto647N. Orange arrowhead indicates the stem dendrite of the displayed AcD neuron. Yellow dashed square indicates the AIS. Scale bar is 10 µm. Bottom row: Single plane 2D gSTED image of F-actin in the AIS of nonAcD and AcD neurons (corresponding zoom-ins of yellow dashed square in top row). Scale bar is 1 µm. Red arrowhead indicates F-actin patch. Black dashed line indicates the F-actin rings shown in (**C**). **(C)** Intensity profile of periodic F-actin structures along the longitudinal axis of the AIS in nonAcD and AcD neurons; corresponding to black dashed lines in (**B**) bottom row. **(D)** Quantification of distance between F-actin rings at the AIS of nonAcD and AcD neurons. Mean ± SEM, 3 independent cultures, nonAcD: n= 45 profiles from 12 cells, AcD: n= 67 profiles from 18 cells.

### The AIS of AcD neurons regulates neuronal polarity by selective permissiveness for axonal cargoes

The AIS helps the axon to retain distinct protein composition by selectively blocking MT based trafficking of dendritic cargoes. This protein filtering feature of the AIS depend on the presence of F-actin patches.^20–22^ Our data showed similar F-actin nanoscale organisation between the AIS of AcD and nonAcD neurons, suggesting that the cargo selectivity of the AIS should remain unchanged in AcD neurons. To experimentally prove this, we used two well-established axonal and dendritic markers: pre-synaptic vesicle-associated Ras-related protein 3A (Rab3A) and transferrin receptors (TfRs)^35,36^. We infected neurons with recombinant adeno-associated virus (rAAV) expressing Rab3A tagged with EGFP (EGFP-Rab3A) to visualise axonal cargoes. For dendritic cargoes, we live-labelled endogenous TfRs of neurons with the fluorescently conjugated ligand Transferrin-Alexa568. The AIS was live-labelled with an anti-NF-CF640R antibody (**Figure S1E**). Time-lapse imaging revealed that EGFP-Rab3A vesicles in AcD neurons were specifically directed into the axon (**Figure 4A-4C**; **Video S8**). Conversely, the TfR vesicles in AcD neurons were halted at the AIS and were mostly moving within somato-dendritic compartments (**Figure 4J-4L**; **Video S10**). These data suggest that, alike the AIS of nonAcD neurons, the AIS of AcD neurons selectively blocks the entry of dendritic cargo into the axonal region.

**Figure 4.**
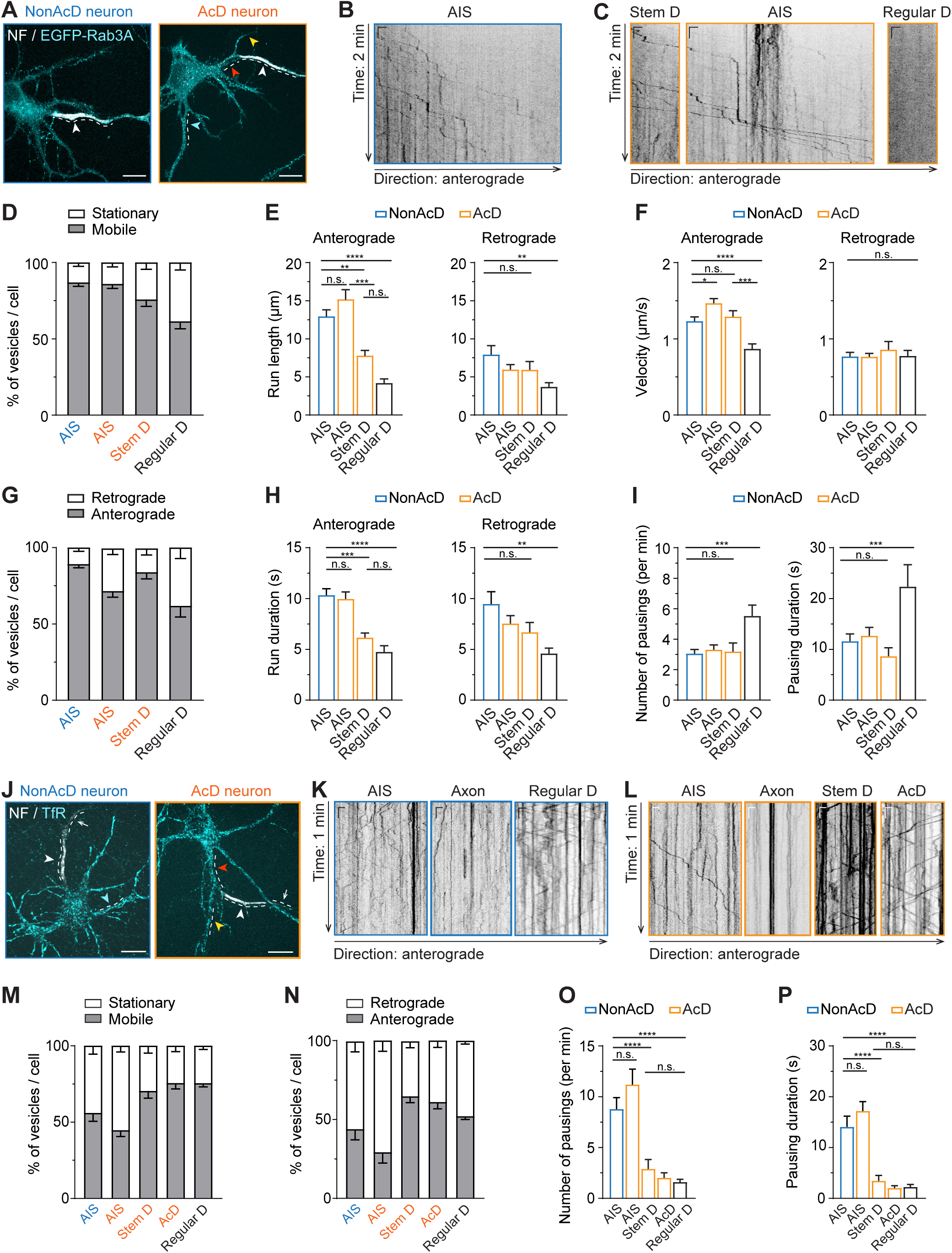
The AIS of AcD neurons serves as an efficient barrier for entry of dendritic cargo into the axon. **(A)** Representative images of nonAcD and AcD neurons expressing a pre-synaptic vesicle marker EGFP-Rab3A, the AIS is live-labelled anti-NF-CF640R (NF). White dashed line indicates the analysed area; white arrowhead indicates the AIS; red, yellow and cyan arrowheads indicate the stem dendrite, AcD and regular dendrite of AcD neuron, respectively. Scale bar is 15 µm. **(B)** Representative kymographs showing trajectories of EGFP-Rab3A vesicles entering the AIS of nonAcD neuron shown in (A), corresponding to area indicated by white dashed line. Scale is 10 s (vertical) and 2 µm (horizontal). **(C)** Representative kymographs showing trajectories of EGFP-Rab3A vesicles moving in the stem dendrite (Stem D), AIS and regular dendrite (Regular D) of AcD neuron shown in (A), corresponding to area indicated by white dashed line. Scale is 10 s (vertical) and 2 µm (horizontal). Anterograde and retrograde direction is the same as in (B). (**D** and **G**) Motility and directionalities of EGFP-Rab3A vesicles in the AIS of nonAcD neuron, the AIS and stem dendrite (Stem D) of AcD neuron, and the regular dendrite (Regular D) of both nonAcD and AcD neurons. (**D**) Percentage of running and stationary EGFP-Rab3A vesicles. (**G**) Percentage of EGFP-Rab3A vesicles running towards anterograde and retrograde directions. Mean ± SEM, 7 independent cultures, AIS (nonAcD): n = 42 cells, AIS (AcD) n = 42 cells, Stem D: n= 28 cells, Regular D: n= 27 cells. (**E**, **F** and **H**) Average length (**E**), duration (**H**) and velocity (**F**) of EGFP-Rab3A vesicles running towards anterograde and retrograde direction within the AIS of nonAcD neuron, the AIS and stem dendrite (Stem D) of AcD neuron, and the regular dendrite (Regular D) of both nonAcD and AcD neurons. Mean ± SEM, 7 independent cultures, Anterograde: AIS (nonAcD) n = 42 cells, AIS (AcD) n = 41 cells, Stem D n= 28 cells, Regular D n= 26 cells, Retrograde: AIS (nonAcD) n = 25 cells, AIS (AcD) n = 37 cells, Stem D n= 17 cells, Regular D n= 20 cells. **(I)** Average number of pauses (left) and total pausing duration (right) of EGFP-Rab3A vesicles moving within the indicated regions of nonAcD and AcD neuron regardless of directions. Mean ± SEM, 7 independent cultures, AIS (nonAcD) n = 42 cells, AIS (AcD) n = 42 cells, Stem D n= 28 cells, Regular D n= 27 cells. **(J)** Representative images of nonAcD and AcD neurons labelled with anti-NF-CF640R (NF) for AIS and Transferrin-Alexa568 for TfRs as dendritic cargo. White dashed line indicates analysed area; white arrowhead indicates the AIS; cyan arrowhead indicates regular dendrite of nonAcD neuron; yellow and red arrowhead indicates the AcD and stem dendrite of AcD neuron, respectively. Scale bar is 15 µm. **(K)** Representative kymographs showing trajectories of TfR vesicles entering and moving in the AIS, axon and regular dendrite (Regular D) of nonAcD neuron shown in (A), corresponding to the area indicated by white dashed line. Scale is 5 s (vertical) and 2 µm (horizontal). Anterograde direction is the same as in (B). (L) Representative kymographs showing trajectories of TfR vesicles moving in the AIS, axon, stem dendrite (Stem D) and AcD of AcD neuron shown in (**A**), corresponding to the area indicated by white dashed line. Scale is 5 s (vertical) and 2 µm (horizontal). Anterograde and retrograde direction is the same as in (**B**). (**M** and **N**) Motility and directionalities of TfR vesicles in the AIS of nonAcD neuron, the AIS, stem dendrite (Stem D) and AcD of AcD neuron, and the regular dendrite (Regular D) of both nonAcD and AcD neurons. (**M**) Percentage of running and stationary TfR vesicles. (**N**) Percentage of TfR vesicles running towards anterograde and retrograde direction. Mean ± SEM, 5 independent cultures, AIS (nonAcD) n = 25 cells, AIS (AcD) n = 23 cells, Stem D n= 23 cells, AcD n = 19 cells, Regular D n= 56 cells. (**O** and **P**) Average number of pause (**O**) and total pausing duration (**P**) of TfR vesicles travelling in the indicated regions of nonAcD and AcD neuron regardless of directions. Mean ± SEM, 5 independent cultures, AIS (nonAcD) n = 25 cells, AIS (AcD) n = 23 cells, Stem D n= 23 cells, AcD n = 19 cells, Regular D n= 56 cells. One-way ANOVA with Tukey’s multiple comparisons test, no significance (n.s.) p > 0.05, **p < 0.01, ***p < 0.001, ****p < 0.0001.

To further assess the axonal cargo permissiveness and dendritic cargo filtering capacity of the AIS in AcD neurons in more detail, we thoroughly analysed the trajectories of EGFP-Rab3A and TfR vesicles using the Python-based program KYMOA (see methods for details). Quantitative analysis showed that the majority of EGFP-Rab3A vesicles were mobile within the AIS of both AcD and nonAcD neurons (nonAcD: 90% mobile, 10% stationary; AcD: 89% mobile, 11% stationary; **Figure 4D**), and no significant difference was found either on the average number or percentage of mobile vesicles between the two cell types (**Figure 4D**; **Figure S2A**; **Video S8** and **S9**). These mobile EGFP-Rab3A vesicles mainly travelled anterogradely across the AIS towards the distal axon (nonAcD: 80% anterograde, 20% retrograde; AcD: 70% anterograde, 30% retrograde; **Figure 4G**) and spent nearly 70% of the total travelling time in processive runs (**Figure S2B**). Only a small portion of EGFP-Rab3A vesicles (nonAcD neuron: 3%; AcD neuron: 5%; **Figure S2C**) underwent on average one direction change while traveling anterogradely through the AIS (**Figure S2C**). During the rest of the total travelling time, the anterogradely transported EGFP-Rab3A vesicles in the AIS of both AcD and nonAcD neurons were either passively diffusing back and forth (**Figure S2B**) or shortly pausing (**Figure S2B**) with on average three pauses per minute, 3 s mean duration of each pause and 10 s of total pausing time (**Figure 4I**; **Figure S2D**). We found that the anterogradely transported EGFP-Rab3A vesicles in the AIS of AcD neurons show slightly longer mean run length, similar mean run duration and, therefore, higher mean run speed than in the AIS of nonAcD neurons (**Figure 4E**, **4H** and **4F**). This finding suggests that, in AcD neurons, the axonal cargoes are transported slightly faster across the AIS. Of note, we did not observe any difference on retrogradely transported EGFP-Rab3A vesicles in the AIS of AcD and nonAcD neurons (**Figure 4E**, **4H** and **4F**), and EGFP-Rab3A vesicles in regular dendrites showed very limited mobility and reduced trafficking (**Figure 4E-4I**; **Figure S2C**-**S2D**).

In comparison to axonal cargoes, dendritic TfR-positive vesicles in both AcD and nonAcD neurons were immobile in the AIS but highly mobile in the dendritic region (**Figure 4M**; **Figure S2E** and **S2F**; **Video S10** and **S11**). Although a few moving TfR vesicles were present at the AIS (**Figure 4M**; **Figure S2E**), these vesicles preferentially moved towards the retrograde direction (**Figure 4N**) and showed a higher percentage of pausing, longer pausing duration, and slower velocity than in dendritic regions (**Figure 4O** and **4P**; **Figure S2F-S2K**), indicating that their trafficking is severely impaired at the AIS. Almost no entries from the AIS into the axon were observed (**Figure 4K, 4L**). Taken together, these results suggest that the AIS of AcD neurons fully preserves its selectivity towards dendritic cargoes, and the anterograde transport of axonal cargoes is equally effective and processive as in the AIS of nonAcD neurons.

Intriguingly, we noticed that both axonal cargo EGFP-Rab3A and dendritic cargo TfRs were actively transported at the stem dendrite of AcD neurons (**Figure 4**; **Video S8** and **S10**). The trafficking velocity of EGFP-Rab3A and TfR vesicles highly resembled that observed in the AIS and dendrites (**Figure 4F**; **Figure S2K**), respectively, except that the average run length and run duration of both cargoes were much shorter at the stem dendrite (**Figure 4E** and **4H**; **Figure S2I-S2K**). This reduction of run length and duration is very likely caused by the difference in physical length, since the stem dendrite is in general shorter than the AIS and the dendrites.

### In comparison to nonAcD neurons, AcD neurons contain less voltage-gated sodium channels, COs and inhibitory synapses at the AIS

Next, we focused on voltage-gated sodium channels (VGSCs) and COs at the AIS of AcD neurons, as both components are critical for the AIS to regulate neuronal excitability. Immunostaining of neurons with an antibody against sodium channels (pan-Nav1) and synpo as a marker of COs indeed revealed an enrichment of VGSCs (**Figure 5A** and **5B**) and formation of COs at AcD neuron’s AIS (**Figure 5C**). However, we found that the intensity of the pan-Nav1 staining and the density of synpo clusters at the AIS of AcD neurons are lower than in nonAcD neurons (**Figure 5D** and **5E**). We also measured the size of synpo clusters at the AIS of both cell types and found no difference (**Figure 5E**). Such alterations in composition could impact the generation of APs as well as calcium handling at the AIS. We also analysed the endogenous expression of the AIS-specific ECM protein brevican in DIV21 AcD neurons (**Figure S3A**). The brevican signal in AcD neurons showed the same localisation as in nonAcD neurons to exclusively accumulate at the beginning of the axon (**Figure S3A** and **S3B**), suggesting that AcD neurons also form specific ECM at the AIS.

**Figure 5.**
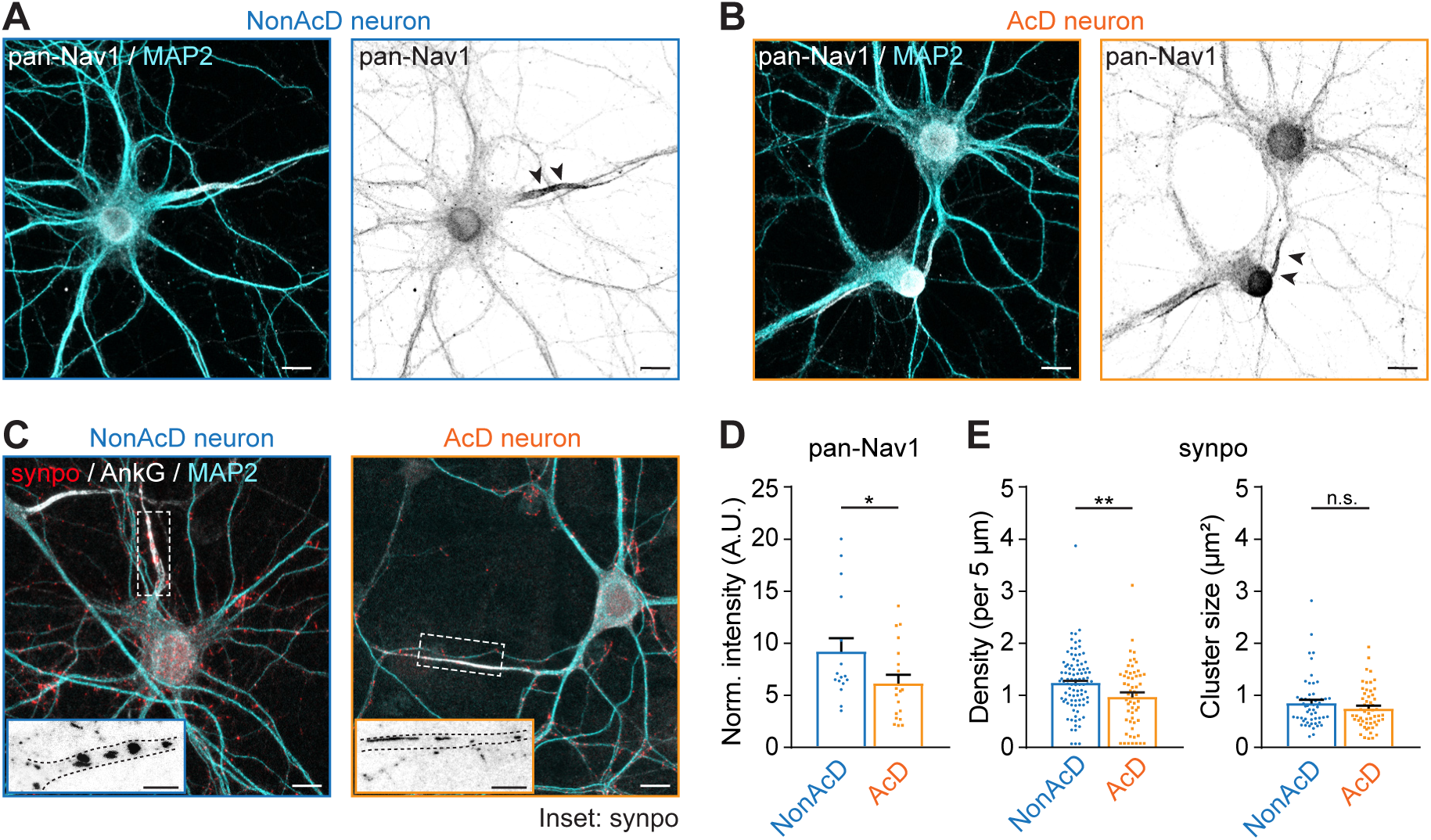
AcD neurons contain less voltage-gated sodium channels and COs than nonAcD neurons at the AIS (**A** and **B**) Representative images of voltage-gated sodium channels labelled by pan-Nav1 staining at the AIS of DIV21 nonAcD (**A**) and AcD (**B**) neurons. Scale bar is 10 µm. (C) Representative images of cisternal organelles (COs) labelled by synaptopodin (synpo) staining at the AIS of DIV21 nonAcD and AcD neurons. Scale bar is 10 µm. Inset: Zoom-ins corresponding to white dashed rectangular. Black dashed lines indicate the edge of AIS. Scale bar is 5 µm. (D) Quantification of pan-Nav1 fluorescent intensity at the AIS of nonAcD and AcD neurons. Mean ± SEM, 3 independent cultures, nonAcD: n= 16 cells, AcD: n= 19 cells. (E) Quantification of synpo cluster density (left) and size (right) at the AIS of nonAcD and AcD neurons. Mean ± SEM, 3 independent cultures, nonAcD: n= 92 cells, AcD: n= 57 cells. Mann-Whitney test: not significant (n.s.) p > 0.05, *p < 0.05, **p < 0.01.

Excitatory neurons are known to establish inhibitory, but not excitatory, connections at the AIS for adjustment of AP initiation.^29^ Therefore, we next set out to quantify the amount of inhibitory synapses at the AIS of AcD neurons. We analysed the density of pre-and post-inhibitory synaptic markers, vesicular GABA transporter (VGAT) and gephyrin, at the AIS of both cell type. We found that the density of pre-and post-synaptic markers at AcD neuron’s AIS are lower than that in nonAcD neurons (**Figure 6A and 6C**), suggesting that AcD neurons receive less inhibitory inputs at the AIS. As a control, we also measured the density of excitatory synapses at the AIS of AcD and nonAcD neurons by using vesicular glutamate transporter (VGLUT) and homer-1 as pre-and post-synaptic markers, respectively. The quantification showed a very low density of both homer-1 and VGLUT at the AIS regardless of axon origin (**Figure 6B** and **6D**), indicating there is no difference in excitatory synapse number.

**Figure 6.**
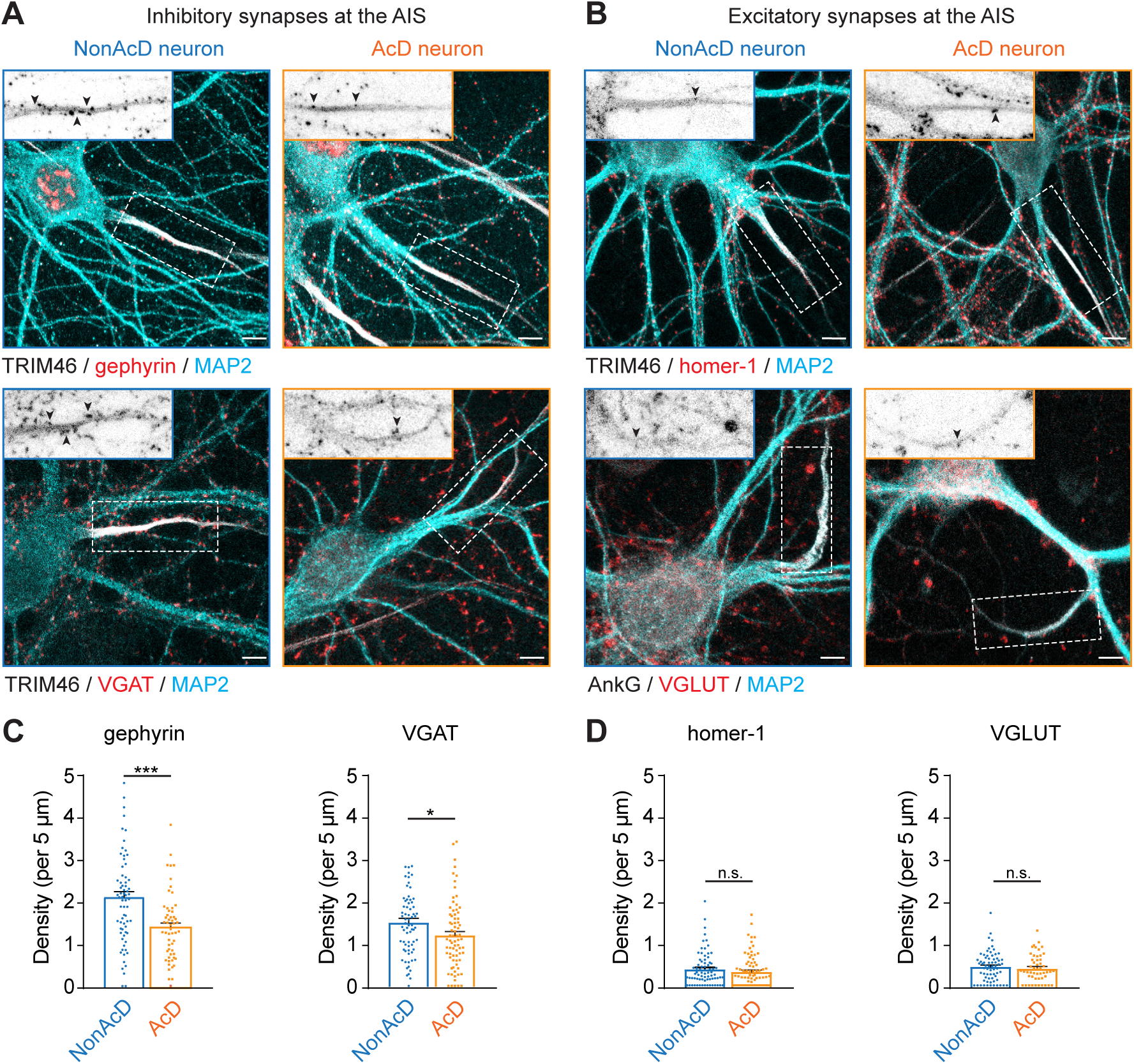
AcD neurons have less inhibitory synapses at the AIS. **(A)** Representative images of nonAcD and AcD neurons stained with markers for pre-and post-synaptic compartments of an inhibitory synapse (VGAT and gephyrin, respectively), for AIS (TRIM46), and for somato-dendritic compartment (MAP2). Upper row: Gephyrin. Lower row: VGAT. Scale bar is 10 µm. Inset: corresponding zoom-ins of white dashed rectangular. Scale bar is 5 µm. Black arrowheads indicate corresponding marker for pre-and post-inhibitory synapse. **(B)** Representative images nonAcD and AcD neurons stained with markers for pre-and post-synaptic compartments of an excitatory synapse (VGLUT and homer-1, respectively), for AIS (TRIM46 or AnkG), and for somato-dendritic compartment (MAP2). Upper row: Homer-1. Lower row: VGLUT. Scale bar is 10 µm. Inset: corresponding zoom-ins of white dashed rectangular. Scale bar is 5 µm. Black arrowheads indicate corresponding marker for pre-and post-excitatory synapse. **(C)** Number of gephyrin and VGAT puncta per 5 µm at the AIS of nonAcD and AcD neurons. Mean ± SEM, 4 independent cultures, gephyrin: n (nonAcD) = 71 cells, n (AcD) = 56 cells, VGAT: n (nonAcD) = 70 cells, n (AcD) = 75 cells. **(D)** Number of Homer-1 and VGLUT puncta per 5 µm at the AIS of nonAcD and AcD neurons. Mean ± SEM, 3 independent cultures for Homer-1, 4 independent experiments for VGLUT, Homer-1: n (nonAcD) = 74 cells, n (AcD) = 78 cells, VGLUT: n (nonAcD) = 67 cells, n (AcD) = 50 cells. Mann-Whitney test: not significant (n.s.) p > 0.05, *p < 0.05, ***p < 0.0001.

### The AIS of AcD neurons does not undergo activity dependent plasticity

The AIS has been shown to regulate neuronal homeostasis at different time scales via two types of plasticity: rapid and chronic plasticity.^27,28^ The rapid plasticity occurs when increasing neuronal activity for a few hours and induces AIS length shortening.^27^ The chronic plasticity on the other hand requires high neuronal activity for at least two days and causes the AIS to shift towards the distal part of the axon without a change in length.^28^ Previous studies showed that the dendritic origin of the axon resulted in different excitability patterns of AcD neurons.^3,4^ This motivated us to investigate whether the AIS of AcD neurons would regulate cellular homeostasis in a different manner. The elevated extracellular K^+^ concentration is an effective way to globally depolarise neurons.^37^ We added KCl into the conditioned medium of DIV12 neurons to elevate extracellular K^+^ concentrations to 15 mM and incubated them for 3 and 48 hours to induce rapid^27^ and chronic AIS plasticity^28^, respectively (**Figure S4A**). To rule out possible effects caused by the change in osmolality, we replaced KCl with NaCl in control groups (**Figure S4A**). We also silenced the baseline synaptic activity of dissociated neurons with 1 µM tetrodotoxin (TTX) as additional control (**Figure S4A**). As read out of AIS plasticity, we stained the AIS with AnkG and measured its fluorescent intensity to define the AIS length and AIS distance (**Figure S4B** and **S4C**). Using a previously published method^27,28^, we considered the start and the end of AIS as the location where AnkG fluorescence intensity drops below 40% of the maximum (**Figure S4C**). The AIS distance is measured as the distance from the start of the axon to the beginning of the AIS (**Figure S4C**).

In nonAcD neurons, we indeed observed a distal shift of the AIS after 48 hours of KCl treatment (**Figure 7A** and **7B**), meaning that the increased extracellular K+ concentration successfully triggered an AIS plasticity. However, we did not observe the previously reported AIS length reduction after 3 hours of KCl treatment (**Figure 7A** and **7B**; **Figure S4D**). Strikingly, in AcD neurons, neither the 48 hours KCl treatment shifted the AIS towards the distal axon tip, nor was the AIS length reduced after 3 hours KCl treatment (**Figure 7C** and **7D**), suggesting that the AIS in AcD neurons is insensitive to elevated activity levels. We also measured the length of the stem dendrite in AcD neurons after 48 hours incubation with KCl, but did not find any difference compared to controls (**Figure S4G**). This indicates that the stem dendrite of AcD neurons does not compensate for abnormal activities as speculated in a previous study.^4^

**Figure 7.**
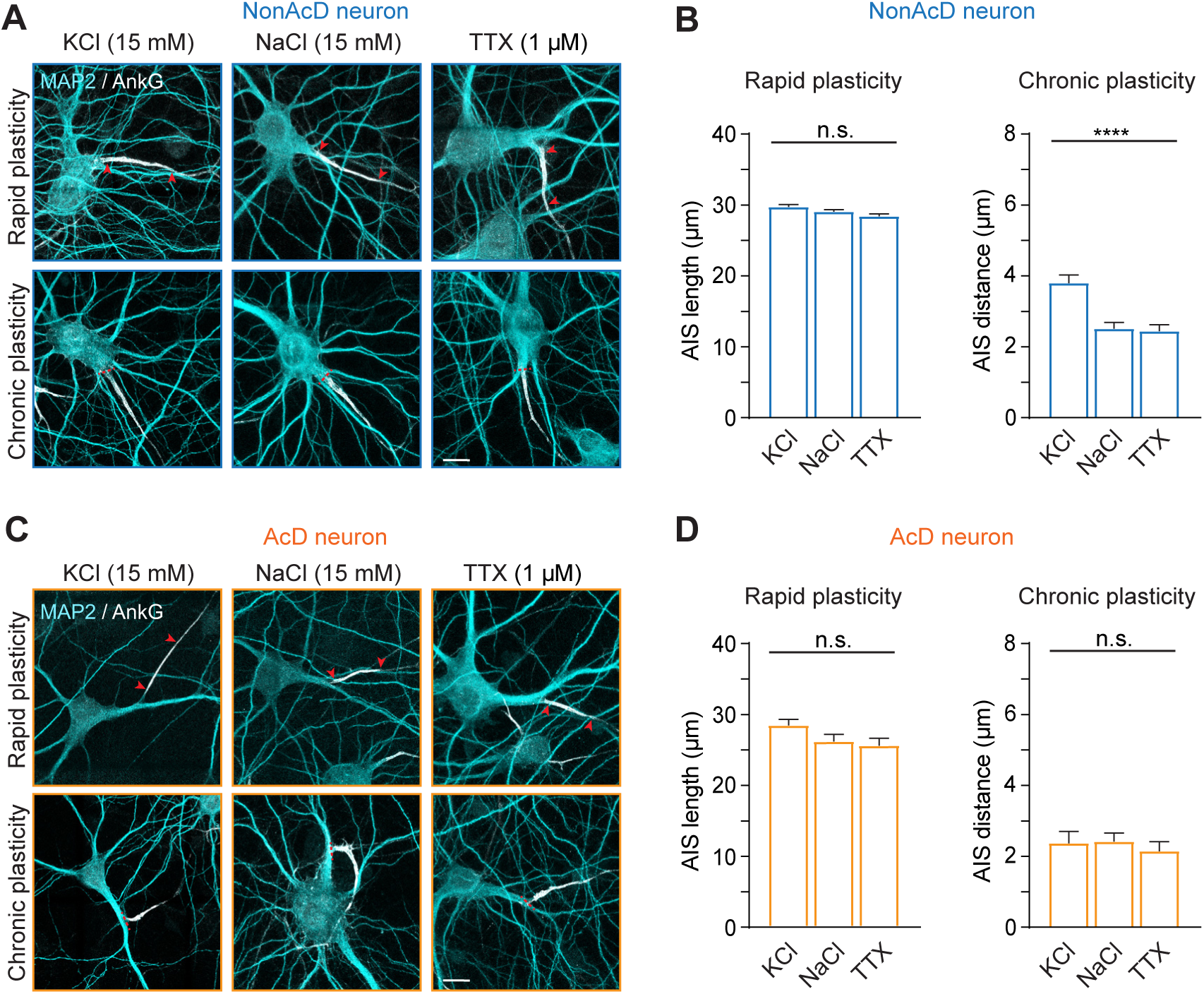
Rapid and chronic plasticity of the AIS in AcD neurons. **(A)** Representative images of the AIS in nonAcD neurons upon induction of rapid (upper row) and chronic (bottom row) plasticity. Neurons were treated with KCl for depolarization, NaCl as control and TTX for silencing of activity. Red arrowheads in upper row indicate the start and end of AIS. Red dashed line in bottom row indicates the start of the axon. Scale bar is 10 µm. **(B)** Measurement of AIS length and AIS distance in nonAcD neurons treated with KCl (depolarization), NaCl (control) and TTX (silencing) for 3 and 48 hours. Mean ± SEM, 3 independent cultures, Rapid plasticity: n (KCl) = 298 cells, n (NaCl) = 264 cells, n (TTX) = 306 cells, Chronic plasticity: n (KCl) = 189 cells, n (NaCl) = 192 cells, n (TTX) = 202 cells. **(C)** Representative images of the AIS in AcD neurons upon induction of rapid (upper row) and chronic (bottom row) plasticity. Neurons were treated with KCl for depolarization, NaCl as control and TTX for silencing. Red arrowheads in upper row indicate the start and end of AIS. Red dashed line in bottom row indicates the start of the axon. Scale bar is 10 µm. **(D)** Measurement of AIS length and AIS distance in AcD neurons treated with KCl (depolarization), NaCl (control) and TTX (silencing) for 3 and 48 hours. Mean ± SEM, 4 independent cultures, Rapid plasticity: n (KCl) = 48 cells, n (NaCl) = 40 cells, n (TTX) = 41 cells, Chronic plasticity: n (KCl) = 73 cells, n (NaCl) = 75 cells, n (TTX) = 66 cells. One-way ANOVA with Tukey’s multiple comparisons test, no significance (n.s.) p > 0.05, ****p < 0.0001.

As reported previously, both NaCl and TTX showed no effect either on AIS length after 3 hours (**Figure 7**), or on AIS position after 48 hours in both AcD and nonAcD neurons (**Figure 7**). ^27,28^ However, we noticed that after 48 hours incubation, TTX specifically increased AIS length in nonAcD neurons (**Figure S4E**), whereas the length of AIS in AcD neurons was unchanged (**Figure S4F**). This finding supports the fact that the AIS also undergoes structural remodelling when nonAcD neurons encounter extremely low activity levels. More importantly, it further indicates that the AIS of AcD neurons is insensitive to activity changes.

To answer the question of what prevents the AIS from undergoing plasticity in AcD neurons, we proceeded to explore possible mechanisms governing the AIS plasticity. At first, we thought that the AIS-specific ECM could cross-link membrane components^38^ and therefore restrict the ability of the AIS to change its size and position in response to neuronal activity. To test this, we investigated the expression of the AIS-specific ECM protein brevican in DIV12 neurons, which is the age of neurons when AIS plasticity was induced in our study. However, we found almost no brevican signal in DIV12 neurons (**Figure S5**), suggesting that the ECM is unlikely to be involved at this time point.

The AIS plasticity in excitatory neurons is triggered by Ca^2+^ signalling.^28^ A previous study already hinted a potential role of COs in rapid AIS plasticity.^39^ We therefore wondered if COs are also changed upon induction of chronic AIS plasticity. To test this, we assessed the density, size and distribution of COs in the AIS of nonAcD neurons treated with KCl for 48 hours. As a control, we first analysed untreated neurons at the same age that chronic AIS plasticity was induced (DIV12 and DIV14). We found that COs were already formed at the AIS of DIV12 neurons (**Figure S6A**). Quantification showed that both the amount and the size of synpo clusters are the same between DIV12 and DIV14 neurons (**Figure S6A-S6C**), suggesting there are no developmental changes of the COs during the time of plasticity induction in our study.

Next, we did the same analysis of COs in neurons treated with KCl for 48 hours to induce chronic AIS plasticity (**Figure S6D** and **S6E**) and compared them to control group. We found that in KCl treated neurons, both the density and number of synpo clusters at the AIS are reduced (**Figure S6F**). We also compared the size of synpo clusters and their distribution within the AIS, but found no difference on both parameters (**Figure S6G** and **S6H**). Collectively, our data suggest that chronically and sustainably increase neuronal activity indeed reduce the amount of COs at the AIS. This result resembles the situation in AcD neurons, which have less COs (**Figure 5**) at the AIS and possibly higher activity rate^3,4^.

## Discussion

AcD neurons have been widely observed across species ranging from invertebrates to vertebrates.^1–4,40^ Only recently, the abundance of AcD neurons in certain brain areas was characterised and their role in memory consolidation was discovered.^2–4^ Despite their importance in brain functions, AcD neurons remain largely unexplored from a cell biology perspective. In this study, we show that the development of AcD neurons is intrinsically driven. We found that in most cases a single precursor neurite gives rise to both the axon and the AcD. Although multiple strategies exist for axon and AcD differentiation, our data clearly indicate that AcD neurons still follow the general developmental sequence^5^ to first establish the axon and then the dendrites, and the AcD is derived mainly from a collateral formed at proximal immature axon during early development. The AIS of AcD neurons is positive for classical AIS markers, it associates with the AIS-specific ECM and has similar F-actin and MT organisation as in nonAcD neurons. Live imaging experiments with axon-and dendrite-specific cargoes demonstrated that the AIS of AcD neurons likewise acts as an efficient filter to selectively block the entry of dendritic proteins into the axon. Interestingly, our data indicate that AcD neurons have fewer sodium channels and inhibitory synapses at the AIS compared to nonAcD neurons. Notably, they also contain fewer COs, resembling the situation in chronically active nonAcD neurons. These differences in the structural characteristics of the AIS suggest that the functionality of the AIS in regulating the excitability and homeostasis of AcD neurons might differ from that of nonAcD neurons. Indeed, we find that the AIS emerging from the dendrite does not undergo homeostatic plasticity to compensate for increased activity levels. Collectively, our data provide new insights into the development of AcD neurons, and demonstrate that there are differences in AIS functionality between AcD and nonAcD neurons.

Our study triggers several exciting new questions. For example, how do AcD neurons specifically form a branch at the proximal part of the precursor axon, stabilise it and then transform as AcD? Or is it rather a stochastic process or a genetically encoded program leading to the development of a specific type of glutamatergic neuron? To answer these questions, it will be important to combine live imaging with interrogation of molecular mechanisms which have been reported to assist neurons in collateral genesis and the fate definition of neurites. Most of these mechanisms involve actin and MT dynamics. For instance, the actin binding protein Arp2/3 relaxes the rigidity of the actin skeleton to promote fast outgrowth of precursor axon^41^, and MT arrays in the precursor axon flow retrogradely with lower speed than in other neurites to locally increase the amount of MTs for protein delivery.^42,43^ In addition, several intrinsic polarity effectors, such as Shootin1^44^, Fmn2^7^ and TRIM46^15,45^, have also been found to participate in neuron development. Thus, future efforts should focus on finding specific proteins and associated signalling pathways that instruct AcD neuron morphogenesis. It might be that transformation into AcD type neurons is a fully stochastic process which depends on probabilities and speed of neurite growth and retraction. In depth analysis of longitudinal imaging data combined with mathematical modelling could help to address this point. It is also possible that the ability to grow an axon from a dendrite comes from AcD neurons being a different subtype of pyramidal neurons. It would be of further interest to perform single cell RNA sequencing to analyse the gene expression profiles of AcD neurons and possibly identify specific markers of this cell type.

The compartmentalization of axonal and dendritic proteins is an essential step in the establishment and maintenance of neuronal polarity. Such segregation is achieved by means of polarized cargo trafficking, which is mediated by the coordinated interplay of cargo adapters, motor proteins, the cytoskeleton, and the presence of a specialized membrane compartment - the AIS. By demonstrating that AcD neurons preserve axonal MT arrangement at the stem dendrite and that their AIS is well-suited for filtering out dendritic cargos, we show how AcD neurons can achieve targeted protein delivery even in the case of an axon starting from a dendrite.

Patch clamp recordings of nonAcD hippocampal excitatory neurons demonstrated that the rapid and chronic plasticity of the AIS fine tunes neuronal excitability to prevent neurons from becoming epileptic.^27,28^ Our results show that the AIS in excitatory AcD neurons does not undergo any structural plasticity upon changes of neuronal activity, suggesting that a different strategy might be employed by AcD neurons to maintain cellular homeostasis. It could also be that AcD neurons are less adaptive than nonAcD neurons which makes them more susceptible to pathological conditions. Therefore, it is of interest to investigate the role of AcD neurons in pathological models related to brain diseases, such as epilepsy or Alzheimer’s disease.

The regulatory mechanisms underlying AIS plasticity are still ambiguous. Up to date, several lines of evidence suggest the significance of Ca^2+^ signalling in this process. The activity of the Ca^2+^-and calmodulin-dependent protein phosphatase calcineurin has been reported to be relevant for AIS plasticity^46^, and specifically blocking voltage gated Ca^2+^ channels (VGCCs) disrupted the distal shift of AIS in hippocampal excitatory neurons^28^. Intracellular calcium stores in form of the CO are particularly interesting in this regard ^23–25^, First, a recent study reported that the number of COs is decreased when rapid AIS plasticity in rat hippocampal DGCs is induced.^39^ Here we could show that upon induction of chronic plasticity, there are less COs in the AIS of nonAcD neurons. This correlates with the fewer COs, we found in the AIS of AcD cells. As the latter ones are presumably more excitable when the input comes from the AcD, it is possible that they have already achieved their plasticity limit in terms of AIS length and localization. Our results suggest that the CO number is more likely to be the consequence than the cause of diminished AIS plasticity in AcD neurons. In the future, one should more thoroughly investigate the mechanistic role of COs in the process of AIS plasticity but also investigate other possible mechanisms, such as the role of sodium channel endocytosis^47^ or the presence and impact of other AIS components which were not explored in this study. Recently, several new AIS specific proteins were discovered via proximity labelling assays.^45^ It would be interesting to test whether they are involved in regulating AIS plasticity and to compare between AcD and nonAcD neurons. Another direction to explore is the link between the AIS-specific ECM and AIS plasticity. Regardless of our data showing the absence of brevican at the age of neuronal cultures when we performed our experiments (DIV12), it is still possible that the ECM plays a role in older neurons, as AIS is plastic throughout the entire lifespan^28,39,48,49^.

Surprisingly, we did not observe any rapid AIS plasticity effect in nonAcD neurons. This might be explained by the fact that hippocampal primary cultures are a mix of different cell types, and a previous study showed that rapid AIS plasticity occurs mainly in hippocampal dentate granule cells (DGCs) but not in principal excitatory neurons.^27^ Another possibility is that certain post-synaptic signalling factors are required to trigger rapid AIS plasticity, as KCl stimulation mainly changes the membrane potential to depolarise neurons but lacks input specificity.^37^ Consistently, a recent study indeed show that the AIS of hippocampal CA1 neurons are shortened after 3 hours incubation with N-methyl-D-aspartate (NMDA) to stimulate NMDA receptors at the post-synapses.^47^ The same protocol of AIS plasticity induction should also be tested in AcD neurons to see whether post-synaptic activities could trigger AIS plasticity.

The inhibitory synapses at the AIS of excitatory neurons is an important modulator for the activation and silencing of specific neuronal circuits.^29^ Our data demonstrate that the AIS of AcD neuron has less inhibitory innervation than in nonAcD neurons, supporting the fact that AcD neurons can evade peri-somatic inhibition during network oscillation. It would be interesting to see in the future which type of interneurons are innervating the AIS of AcD neurons and how this contributes to the higher intrinsic excitability of AcD neurons.

## Acknowledgements

We would like to thank Bianca Slivinschi for experimental assistance and help with data analysis, Zhengyue Zhang (MUNI-CEITEC, Brno) for help with Python scripting, Tomas Fanutza (ZMNH, Hamburg) and Karin Ruban (CSSB, Hamburg) for help with sample preparation and transport between the institutions, Julia Sandberg (CSSB, Hamburg) and Simone Traeger (CSSB, Hamburg) for technical support, Dr. Antonio Virgilio Failla (UMIF, Hamburg) for access and help with use of the STED confocal microscope, and Dr. Christian Conze (LIV, Hamburg) for help with microscope access at LIV. We acknowledge Dr. Anja Konietzny (ZMNH, Hamburg) for technical assistance and valuable feedback on the manuscript, Dr. Ingke Braren (UKE Virus Facility) for rAAV production. This research is supported by the Landesforschungsförderung Hamburg (LFF to MM and KG) and by the German research foundation (Excellence Strategy – EXC-2049–390688087 to MM, DFG FOR5228 to MM and DFG FOR2419 to MM and KG).

## Author contributions

Designed experiments (YH, RT, KG, MM), performed experiments (YH), analysed and interpreted data (YH, KG, MM), developed the manuscript concept (YH, MM), wrote the first draft (YH, MM), reviewed and edited the paper (YH, RT, KG, MM) and acquired funding (KG, MM)

## Competing interests

The authors declare no competing financial interests.

## Materials & Correspondence

Materials (constructs) can be requested from Marina Mikhaylova; correspondence can be addressed to Kay Grünewald and Marina Mikhaylova. Note, the ^2^ Guest Group “Neuronal Protein Transport” at Centre for Molecular Neurobiology has been discontinued in August 2023.

## Methods

### Animals

Wistar Unilever HsdCpb:WU (Envigo) rats were used in this study. Rats were bread and kept at the animal facility of the University Medical Center Hamburg-Eppendorf, UKE, Hamburg, Germany. Animal experiments were carried out in accordance with the European Communities Council Directive (2010/63/EU) and the Animal Welfare Law of the Federal Republic of Germany (Tierschutzgesetz der Bundesrepublik Deutschland, TierSchG) approved by the city-state Hamburg (Behörde für Gesundheit und Verbraucherschutz, Fachbereich Veterinärwesen, from 21.04.2015, ORG781 and 1035) and the animal care committee of the University Medical Center Hamburg-Eppendorf.

### Primary hippocampal neuron preparation and transfections

Primary rat hippocampal neurons were prepared and maintained as described previously^24^. Briefly, hippocampi were extracted from E18 rat embryos and treated with 0.25% trypsin for 15 min at 37°C. Afterwards, hippocampi were physically dissociated by pipetting through a 26G needle and filtered to remove large clumps. The cell suspensions were then plated on poly-l-lysine -coated 18 mm glass coverslips or 35 mm glass bottom petri dish at a density of 20,000 cells (extra low density) or 40,000 cells (low density) or 60,000 cells (high density) per 1 ml in DMEM supplemented with 10% fetal calf serum and antibiotics. After 1 hour, the plating medium was replaced by BrainPhys neuronal medium supplemented with SM1 and 0.5 mM glutamine. Cells were grown at 37°C with 5% CO2, and 95% humidity.

For microtubule orientation experiment, primary neurons were transfected with EB3-Tdtomato at days in vitro (DIV) 14 by using lipofectamine 2000. Conditioned neuronal medium was removed and stored at 37°C with 5% CO2 before transfection. Neurons were then incubated with transfection medium (transfection mixture in BrainPhys medium without supplements) at 37°C with 5% CO2 for 1 hour. After incubation, the transfection medium was exchanged back to conditioned medium, and neurons were imaged 12 hours after. For EGFP-Rab3 trafficking experiment, primary neurons were infected with rAAV9-syn-EGFP-Rab3 virus (final concentration: 1.38E+10 vg/ml) at DIV5 and imaged at DIV7.

### Constructs and recombinant adeno-associated viruses

Detailed information of constructs and virus used in this study can be found in the key resource table. The EB3-Tdtomato construct was obtained from Addgene (Addgene plasmid #50708; http://n2t.net/addgene:50708; RRID:Addgene_50708). For rAAV9-syn-EGFP-Rab3A construct, full length Rab3A sequence was amplified from EGFP-Rab3A vector (Addgene plasmid # 49542) using PCR. The amplified sequence was then cloned onto a recombinant adeno-associated virus (rAAV) backbone with an EGFP tag behind a human synapsin promotor using Cold Fusion cloning kit. The Rab3A sequence is placed behind EGFP sequence for N-terminal tagging of Rab3A. After verification by sequencing, rAAV9 were produced by the UKE virus facility.

### Immunocytochemistry (ICC)

Neurons were fixed in 4% Roti-Histofix (Carl Roth), 4% sucrose in PBS for 10 min at 37°C. The fixation reagent was removed, and coverslips were washed three times with PBS. Subsequently, neurons were permeabilized in 0.2% Triton X-100 in PBS for 10 min, then washed three times in PBS and blocked for 45 min at RT with blocking buffer (BB: 10% horse serum, 0.1% Triton X-100 in PBS). Incubation with primary antibodies was performed in BB at 4°C overnight. After 3× wash in PBS, neurons were incubated with corresponding secondary antibodies in BB for 1.5 h at RT and unbound antibodies were washed out using PBS. In order to distinguish dendrites from axons, pre-conjugated MAP2-Alexa488 antibody diluted in BB was then applied to neurons for 1.5 h at RT. Finally, coverslips were washed three times in PBS (10 min interval), one time in H2O for 10 s and mounted on microscope slides with Mowiol.

### Spinning disc confocal microscopy

For live imaging of EB3-Tdtomato, EGFP-Rab3A and Transferrin receptors (TfRs), spinning-disc confocal microscopy was performed on a Nikon Ti-2E controlled by NIS Elements 5.2 software. Illumination was done by 488 nm, 561 nm, and 642 nm excitation lasers from Omicron laser unit coupled to a Yokogawa CSU-W1 spinning disc unit. Emission was collected through a bandpass filter (Semrock, 442/525/609/700 nm) on an Andor iXon ultra 888 EMCCD camera. Use of 100x TIRF objective (Nikon, ApoTIRF 100x/1.49 oil) achieved a pixel size of 130 nm.

For imaging of neuronal development, phase contrast imaging was performed on a Nikon Ti-2E microscope controlled by NIS Elements 5.2 software. The microscope was equipped with an Omicron laser unit coupled to a Yokogawa Borealis-enhanced CSU-W1 spinning disc unit, transmitted illumination and an Andor iXon ultra 888 EMCCD camera. Samples were illuminated by transmitted light and imaged via 40X objective (Nikon, ApoLWD Lambda 40x/1.15 WI). The final fluorescent map was acquired through the same objective with laser illumination at wavelength of 488 nm and 642 nm. Both microscopes were equipped with a live imaging system from OKO lab and live imaging was performed at 37°C with 5% CO2 and 95% humidity.

### Laser scanning confocal microscopy and STED imaging

Fixed and stained primary hippocampal neurons were imaged at a Leica SP8 confocal microscope (Leica microsystems, Mannheim, Germany). The microscope was controlled by Leica Application Suite X (LASX) software and equipped with a white light laser. Samples were imaged using a 63x oil objective (Leica, 63x HCX PL APO /1.40 oil). Fluorophores were excited at desired wavelength and signals were detected using HyD detectors. Tile scans of maximum 2 mm x 2 mm areas were performed to increase the chance of finding AcD neurons. A single z-stack tile was acquired with dimension of 1024 x 1024 pixels, pixel size of 80 nm, pixel depth of 16 bit, and z-step size of 0.5 µm. Tiles were merged by LASX function Mosac Merge with 10% overlap.

An Abberior gatedSTED system equipped with a 405 to 640 nm pulsed laser and a 60x oil objective (Nikon, P-APO 60x/1.40 oil) was used for confocal and gated STED imaging. For excitation, 640 nm laser was used for Atto647N/Abberior Star 635p, 561 nm laser was used for Abberior Star 580, and 488 nm laser was used for Alexa 488. STED was achieved with a 775 nm pulsed depletion laser for Abberior Star 580, Abberior Star 635p and Atto647N. Emission spectra were collected between 650-720 nm, 605-625 nm and 500-550 nm. Detector time gates were set to 8 ns for all fluorophores. Images were acquired as single planes with a pixel size of 20 x 20 nm (x and y) and 16-bit pixel depth. The corresponding confocal images were acquired with identical settings.

### Classification of AcD and nonACD neurons

AcD and nonAcD neurons were classified based on MAP2 and AnkG stainings which label somato-dendritic compartment and the AIS of axon, respectively. The distance from the starting point of an axon to the adjacent edge of corresponding cell body was referred as axon distance in this study and used as main factor for AcD neuron classification (**Figure 1B**; **Figure S1D**). To measure the axon distance, axon and cell body of a neuron was defined individually based on AnkG and MAP2 signal. A 2 pixel wide segmented line was then drew along the longitudinal axis of stem dendrite to connect the beginning of the axon and the ending edge of the cell body (**Figure 1B**). The length of the segmented line represents axon distance. If the axon origin was not parallel to the ending edge of soma (**Figure S1D**), the axon distance was then considered as the distance from the edge of soma to the axis perpendicularly extended from the centre of the axon origin (**Figure S1D**). Neurons with axon distance larger than 3 µm was consider as AcD neuron. The AcD and nonAcD neuron was classified in the same manner through the entire study, unless otherwise stated.

### Assessment of AcD neuron population in dissociated cultures and the timeline of AIS formation

Dissociated neurons (60,000 cells/ml) were fixed at DIV3, DIV5, DIV7 and DIV12 and immunostained with anti-MAP2-Alexa488 antibody to label the cell body and dendrites. Antibodies against AnkG was used to visualise the AIS. Tile scan of large area was taken for both MAP2 and AnkG channels with 1% laser power and 5 µm depth in z. Total number of neurons at age of DIV5 and DIV7 was counted from all tile scan images. The number of AcD neuron was then counted from the same images and divided by total number of neurons to calculate percentage of AcD neuron population.

### Time-lapse imaging to visualise neuronal development in dissociated culture

High density neurons (60,000 cells/ml) were cultured on a 35 mm glass bottom petri dish with 20 x 20 mm grid coordinates. A 3 mm diameter hole was drilled on the lid of the petri dish for replenishment of conditioned medium. To prevent medium evaporation during the entire imaging period, conditioned medium was refilled on daily basis. Additional three 35 mm petri dishes were filled with distilled H2O and placed in the live imaging chamber without lids to increase atmospheric humidity. The H2O petri dishes was refilled every 12 hours to maintain proper humidity level in the live imaging chamber.

Developmental sequences were recorded 6 hours post plating. Dishes were placed on a spinning disc confocal microscope and imaged for 5 days every 3 hours. To increase the number of AcD neurons, a 7 x 7 mm area was scanned spirally starting from the centre of the grid coordinates. On day 5, neurons were fixed and immunostained with pre-conjugated anti-MAP2-Alexa488 and anti-AnkG antibody to label dendrites and the AIS, respectively. A single plan fluorescent image of both MAP2 and AnkG channels was then taken at the same scanned area with 20% laser power and 200 ms exposure time. The fluorescent image was used as a final map for selection of AcD and nonAcD neurons, and the developmental sequence of selected neurons was retrieved according to grid coordinates. The formation and growth of AcD and axon in AcD neurons was analysed manually by going through the corresponding developmental sequence frame by frame. All analysis was performed on Fiji.

### F-actin staining and analysis of periodic membrane actin cytoskeleton of the AIS

For F-actin staining, extra low density neurons (20,000 cells/ml) were plated on 18 mm high precision glass coverslips, fixed on DIV14, and immunostained with anti-βIV-Spectrin and pre-conjugated anti-MAP2-Alexa488 antibody to label the AIS and dendrites, respectively. Following the antibody staining, the F-actin was labelled by incubation with phalloidin-Atto647N (1:100 dilution in PBS) at 4°C overnight. Following 3x wash with PBS, coverslips were mounted using moviol and imaged on an Abberior gatedSTED system as described above.

Peak Cal 3.0, a custom-written python-based script (see key resource table), was used to measure the F-actin periodicity at the AIS. Briefly, AIS regions with less phalloidin-Atto647N background signal were selected manually from the STED image. A 3 x 3 pixel segmented line profile was drew at the selected regions along the longitudinal axis of the AIS, and the corresponding phalloidin-Atto647N intensity was then extracted using Fiji. The F-actin rings are hence represented by the periodical peaks of phalloidin-Atto647N fluorescent signal along the segmented line. The extracted intensity profile was loaded into Peak Cal 3.0 and smoothed over five pixels to filter the background noise. The indices of phalloidin peaks were then detected using python built-in function Find Peaks. The distance was calculated by subtracting the indices between two adjacent peaks and then times 20 nm pixel size.

### Microtubule extraction and staining

The staining of tyrosinated and acetylated microtubules (MTs) was performed accordingly to previously described protocol^33^. In short, extra low density neurons (20,000 cells/ml) were grown on 18 mm high precision glass coverslips till DIV10. Then neurons were pre-extracted for 1 min using MT-Extraction buffer (0.3% Triton X-100, 0.1% glutaraldehyde, 80 mM PIPES, 1 mM EGTA, 4 mM MgCl2, pH 6.8) and fixed for 10 min using EM grade 4% PFA, 4% sucrose in PBS at 37°C. After 3x wash in PBS, neurons were immunostained with antibodies against tyrosinated and acetylated α-tubulin for dynamic and stable microtubules, respectively. Antibody against TRIM46 was used to locate the AIS. Samples were imaged on an Abberior gatedSTED system as described above. The distribution of stable and dynamic MTs along the width of a region of interest was displayed by extracting and plotting the fluorescent intensities of a 2x2 pixel line along the latitudinal axis.

### Imaging of MT dynamics

High density neurons (60,000 cells/ml) were grown on 35 mm glass bottom petri dish and transfected with EB3-Tdtomato plasmid at the age of DIV13 and imaged 12 hours post transfection using spinning disk confocal microscope. Shortly before imaging, conditioned medium was replaced by BrainPhys medium containing neurofascin-CF640R antibody (1:200 dilution) to label the AIS. Dishes were return in the incubator and kept at 37°C with 5% CO2 for 5 min. Afterwards, conditioned medium was exchanged and neurons were return in the incubator for 10 min to recover. AcD and nonAcD neurons were then identified based on neurofascin and EB3-Tdtomato channel. For that images were acquired as z-stacks with 0.7 µm step size, 5% laser power and 100 ms exposure time. Then, EB3 channel of selected neurons was recorded as a time lapse with frame rate of 1.3 s / frame and 5% laser power for 5 min.

### Axonal and dendritic cargo trafficking assays

For axonal cargo trafficking assay, high density DIV5 neurons (60,000 cells/ml) on 35 mm glass bottom petri dishes were infected for 2 days with rAAV9 virus expressing EGFP-Rab3A. At DIV7, the AIS was labelled at 37°C with 5% CO2 for 5 min by replacing conditioned medium with BrainPhys medium containing neurofascin-CF640R antibody (1:500 dilution). Neurons were then placed back to conditioned medium for recovery and recording. AcD and nonAcD neurons were selected based on EGFP-Rab3A and neurofascin-CF640R fluorescence. Z-stack image was taken for both channels with 0.7 µm step size, 5% laser power and 100 ms exposure time to verify neuronal morphology. To analyse the newly delivered EGFP-Rab3A vesicles and to improve the signal to noise ratio, the vesicles residing at the AIS, stem dendrite and regular dendrite were photo-bleached using 405 nm laser with 70% laser power for 5 s. New EGFP-Rab3A vesicles travelling through the photo-bleached regions were continuously imaged with 5 frames per second and 10% laser power for 2 minutes.

For dendritic cargo trafficking assay, the AIS of high density DIV7 neurons (60,000 cells/ml) grown on 35 mm glass bottom petri dishes was labelled as described above. Then, pre-conjugated Transferrin-Alexa568 (1:1000 dilution) was used to label the endogenous Transferrin receptors (TfRs) by incubation for 15 min at 37°C with 5% CO2 in BrainPhys medium. Then neurons were switched back to the conditioned medium for recovery. AcD and nonAcD neurons were selected based on Transferrin-Alexa568 and neurofascin-CF640R channels. Neuronal morphology was confirmed by z-stack imaging (0.7 µm step size, 5% laser power and 50 ms exposure time) of both channels. Trafficking of TfRs in the AIS, stem dendrite, AcD and regular dendrite was recorded with 5 frames per second and 10% laser power for 2 minutes.

### Analysis of microtubule dynamics and cargo trafficking assays

Time lapse imaging data for EB3-Tdtomato, EGFP-Rab3 and Transferrin-568 were analysed using Fiji and self-written python scripts KYMOA 6.0 and KA Post Processing 2.0 (https://github.com/HU-Berlin-Optobiology/AIS-project.git). In short, segmented lines were drew on time lapse images at region of interests (ROIs). The segmented line is along the anterograde direction (e.g., cell body to distal axon), and the width of the segmented line covers the width of the ROIs to include majority of the signals. Corresponding kymograph were then generated using Fiji plugin KymoResliceWide. Coordinates of each trajectory on the kymograph were extracted using freehand tool and get-coordinate function in Fiji. Subsequently, the coordinates were further processed in KYMOA 6.0 and KA Post Processing 2.0, as described by Konietzny et al. (https://doi.org/10.1101/2023.10.18.562903). For EGFP-Rab3 and Transferrin-568, we defined vesicles having at least one movement, i.e. frame-to-frame change in the y-coordinate, over 5 continuous pixels and frames as mobile vesicles, and the corresponding movement was considered as a run. Otherwise, the vesicles were classified as stationary. Several parameters were then calculated only for mobile vesicles, such as run length, run speed, run duration and net displacement. To define the travelling direction of a mobile vesicle, we took advantage of the net displacement. Mobile vesicles having a displacement above zero were classified as anterogradely transported vesicles, however, below zero were classified as retrogradely transported vesicles. During active transport, vesicles sometimes shortly pause at one spot for a few seconds. We thus defined pausing behaviour as a mobile vesicle that stalled at the same position for at least 3 frames. It is worth noting that we often observed a vesicle repeatedly moving one pixel, stop for one frame and moving for one pixel again. Since those vesicles were neither fully stationary nor being actively transported, we specified this type of movement as passive movement. For EB3-Tdtomato, the running and pausing threshold was set to 2 pixels and 2-10 frames respectively. There was no threshold for passive movement because MT plus ends have unidirectional movements.

### Induction of rapid and chronic AIS plasticity

Induction of AIS plasticity in dissociated primary neuron cultures was performed following previously published protocols.14,15 Briefly, high density dissociated primary neurons (60,000 cells/ml) were depolarised by 15 mM KCl to artificially increase neuronal activity, 15 mM NaCl was used as osmolarity control and 1 µM Tetrodoxin (TTX) was used to silence neuronal activity of the entire culture. For rapid plasticity, neurons were treated for 3 hours at DIV12 and then fixed for 10 min at 37°C. For chronic plasticity, neurons were treated from DIV12 to DIV14 for 48 hours then fixed for 10 min at 37°C. Fixed neurons were immunostained with pre-conjugated anti MAP2-Alexa488 and anti AnkG antibody to label dendrites and AIS respectively and imaged using laser scanning confocal microscope. Tile scan images were taken with 1% laser power for both channels and 5 µm depth in z.

### Analysis of AIS length and distance

The length of AIS and its distance to the starting point of axon was measured based on AnkG fluorescent intensity. Maximum projection of confocal z-stack images was used for this analysis. A 3 pixel wide segmented line was drew from the beginning of the axon towards the distal end to cover the entire AIS. The florescent intensity of AnkG staining along the segmented line profile was extracted in Fiji and processed by AIS Pack 4.0, a self-written python script (https://github.com/HU-Berlin-Optobiology/AIS-project.git). The AnkG intensity profile was smoothed over 4 µm to filter out background noise and the peak of the intensity profile was detected. To define the start and the end of the AIS, the algorithm iterate through intensity values from the peak to the left and right side until the value is below 40% of the peak value. Consequently, the left edge of the AnkG intensity profile is the start of the AIS and the right edge is the end. Indices of the left and right edge are subtracted to calculate AIS length, and the AIS distance equals to the left edge of AnkG intensity profile.

### Immunostaining of inhibitory and excitatory synapses and cisternal organelle

Low density DIV21 neurons (40,000 cells/ml) were fixed for 10 min at 37°C. For inhibitory and excitatory synapse, fixed neurons were stained either with antibody against VGAT and gephyrin for pre-and post-synaptic sites of inhibitory synapse, or with antibody against VGLUT and homer-1 to visualise the pre-and post-synaptic compartments of excitatory synapse. The AIS was stained either with AnkG or TRIM46 antibody depending on antibody species compatibility. MAP2-Alexa488 antibody was used to visualise cell bodies and dendrites. For the CO, the AIS, cell body and dendrite of fixed neurons were stained in the same way as above. The CO was immunostained by antibody against synaptopodin. All samples were imaged using laser scanning confocal microscope, and a tile scan image was taken with 0.5-1% laser power and 5 µm depth in z. The number of synaptopodin puncta as well as number of inhibitory and excitatory synapses were quantified using Fiji. Maximum projection of tile scan images was used for the analysis. Synaptopodin, VGAT, gephyrin, VGLUT and homer-1 puncta within the AIS region were counted using multipoint tool and normalised to the length of measured AIS.

### Immunostaining of AIS specific ECMs and sodium channels

Low density DIV21 neurons (40,000 cells/ml) were fixed for 10 min at 37°C. For AIS-specific ECMs, fixed neurons were stained with pre-conjugated antibody against brevican. For sodium channels, fixed neurons were stained with a polyclonal antibody against alpha subunits of voltage-gated sodium channels (pan-Nav1). MAP2-Alexa488 antibody was used to visualise cell bodies and dendrites. All samples were imaged using laser scanning confocal microscope, and a tile scan image was taken with 5-10% laser power and 5-7 µm depth in z. The florescent intensity of pan-Nav1 was quantified using Fiji. Maximum projection of tile scan images was used for the analysis. Pan-Nav1 intensity was outlined with polygon shape tool. Average intensity of pan-Nav1 was extracted from the outlined shape and normalised to background of each image.

### Analysis of CO size and distribution in the AIS

The CO size was measured using a self-written FIJI plugin synpo_det (https://github.com/HU-Berlin-Optobiology/AIS-project). Briefly, the AIS was manually outlined based on AnkG and MAP2 immunofluorescence using polygon shape tool in FIJI. Single channel image containing Synpo staining as CO marker was duplicated and thresholded. The thresholded image was then processed with find-edges filter to highlight Synpo clusters within the AIS. The area size of the outlined Synpo clusters was then measured to represent Synpo cluster size.

To analyse the distribution of COs within the AIS, a segmented line (3 pixel width) was first drew based on AnkG and MAP2 staining along the central axis of the AIS in FIJI. The line extend from the start of the axon to the distal direction and covers the entire AIS. Subsequently, the coordinates of the pixels along the segmented line was extracted as a reference. Next, the coordinate of each Synpo clusters within the AIS was extracted in FIJI. The Synpo cluster coordinates were then traced back to the reference coordinates extracted from the segmented line using a self-written Python program “AIS_synpo_cluster_analysis”. The distance of each traced Synpo cluster coordinate to the beginning of reference line was then measured to represent the location of COs in the AIS.

### Statistical Analysis and Image representation

Statistical analysis was performed on Prism version 7.03 (GraphPad). Data are represented as percentage or as mean ±SEM. Individual channels in multi-colour images are contrasted for better representation, with set minimum and maximum identical if groups need to be compared. No other modifications were done, unless otherwise stated. All analysis was done on raw images.

## Key resource table

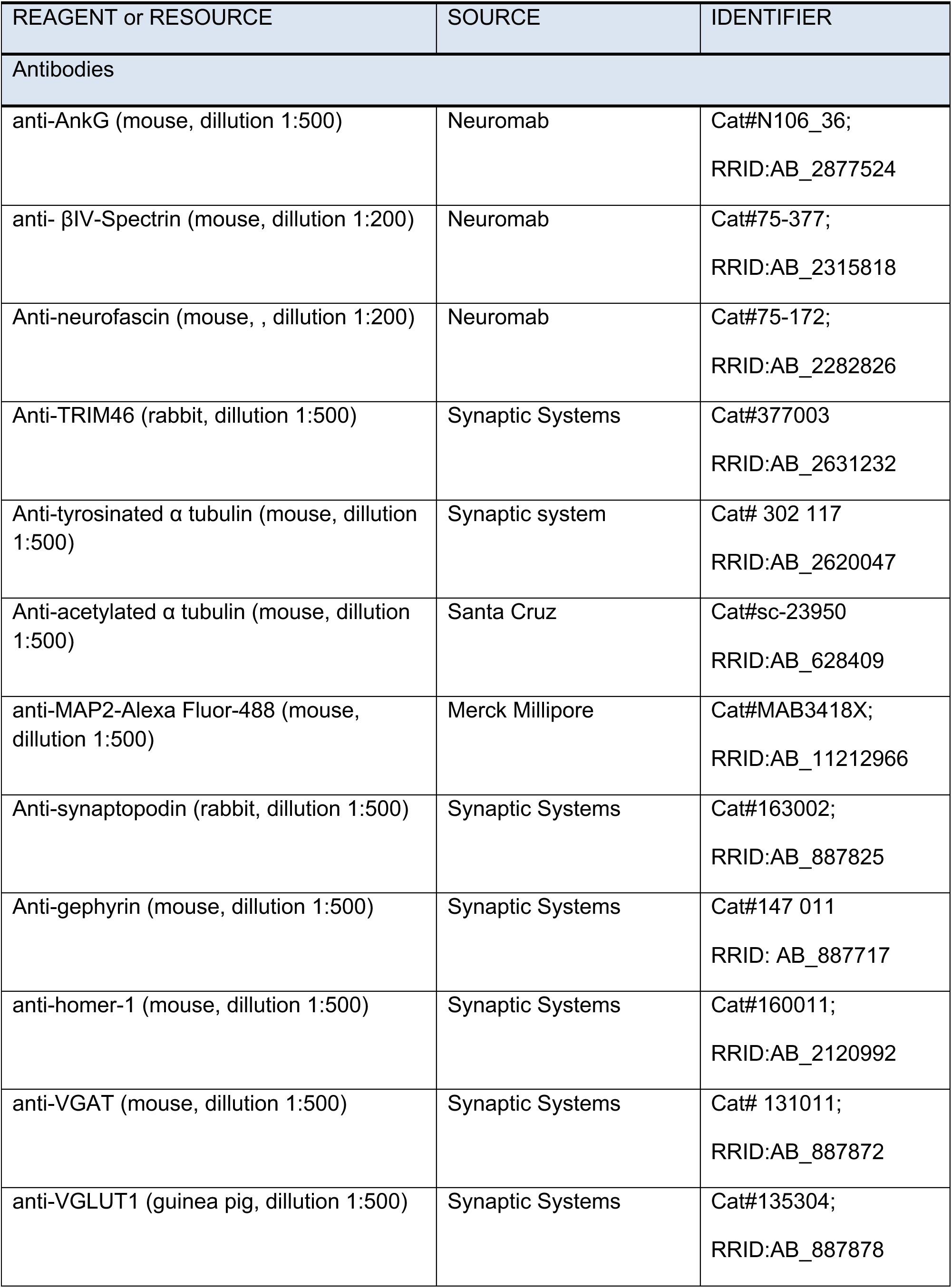
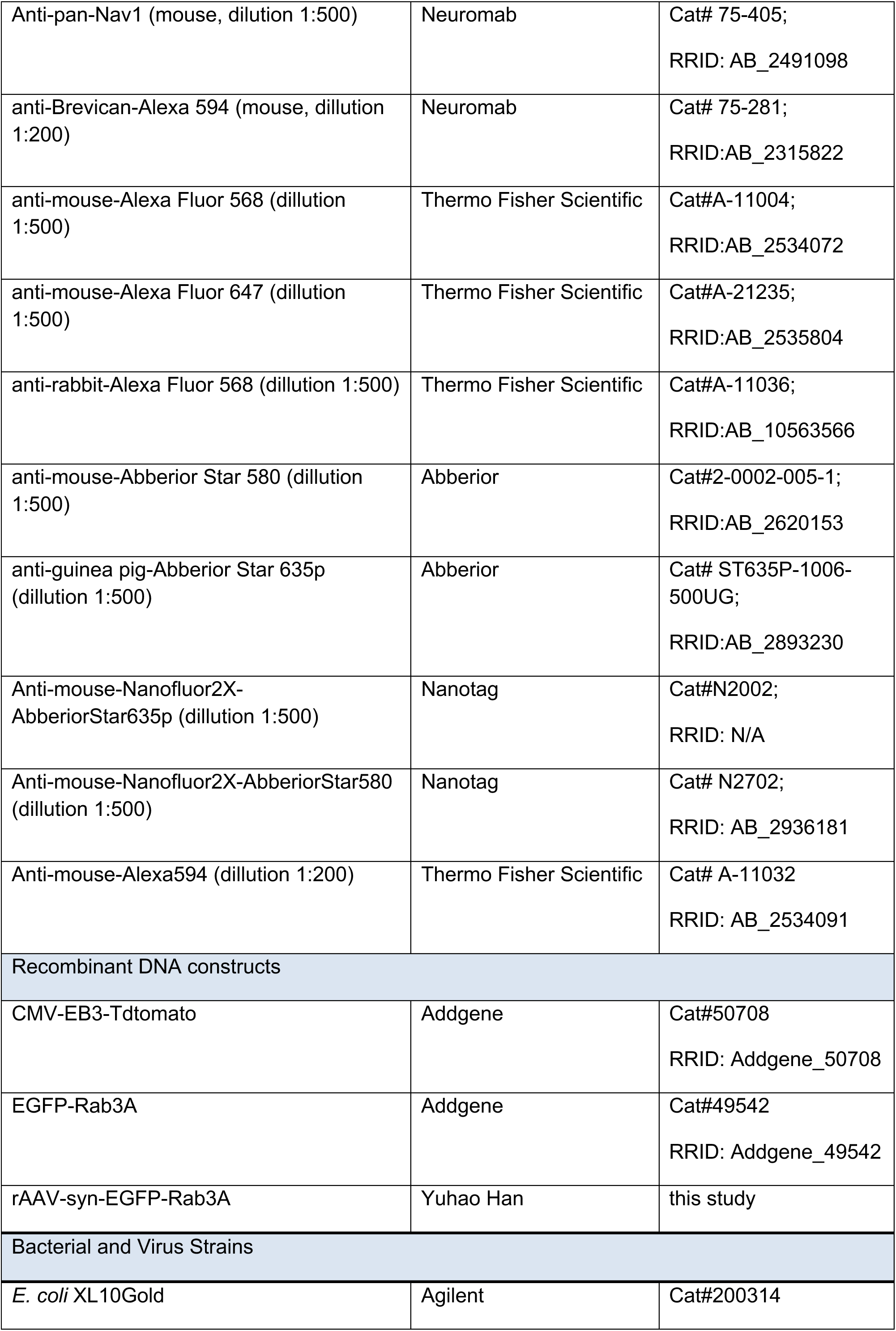
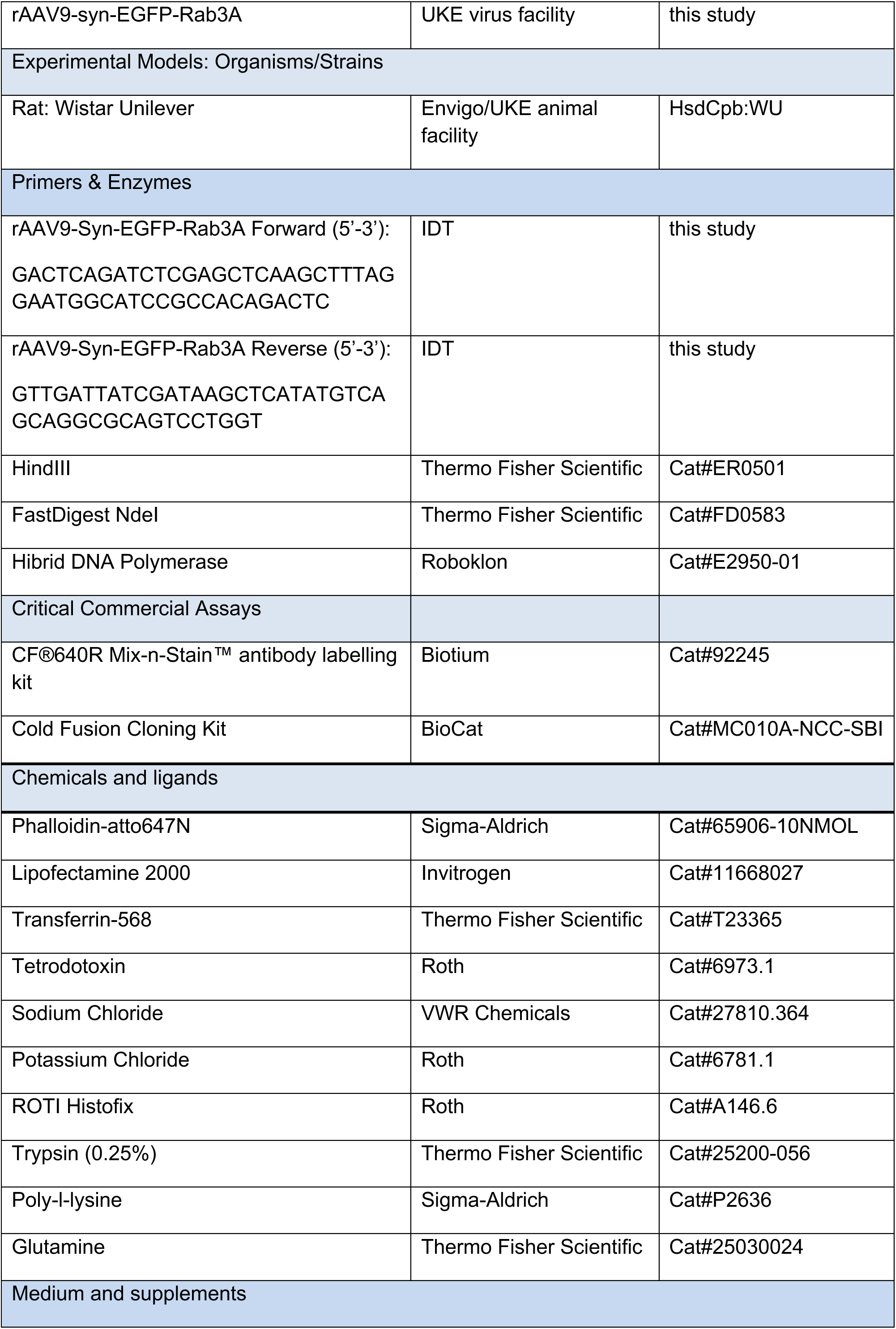
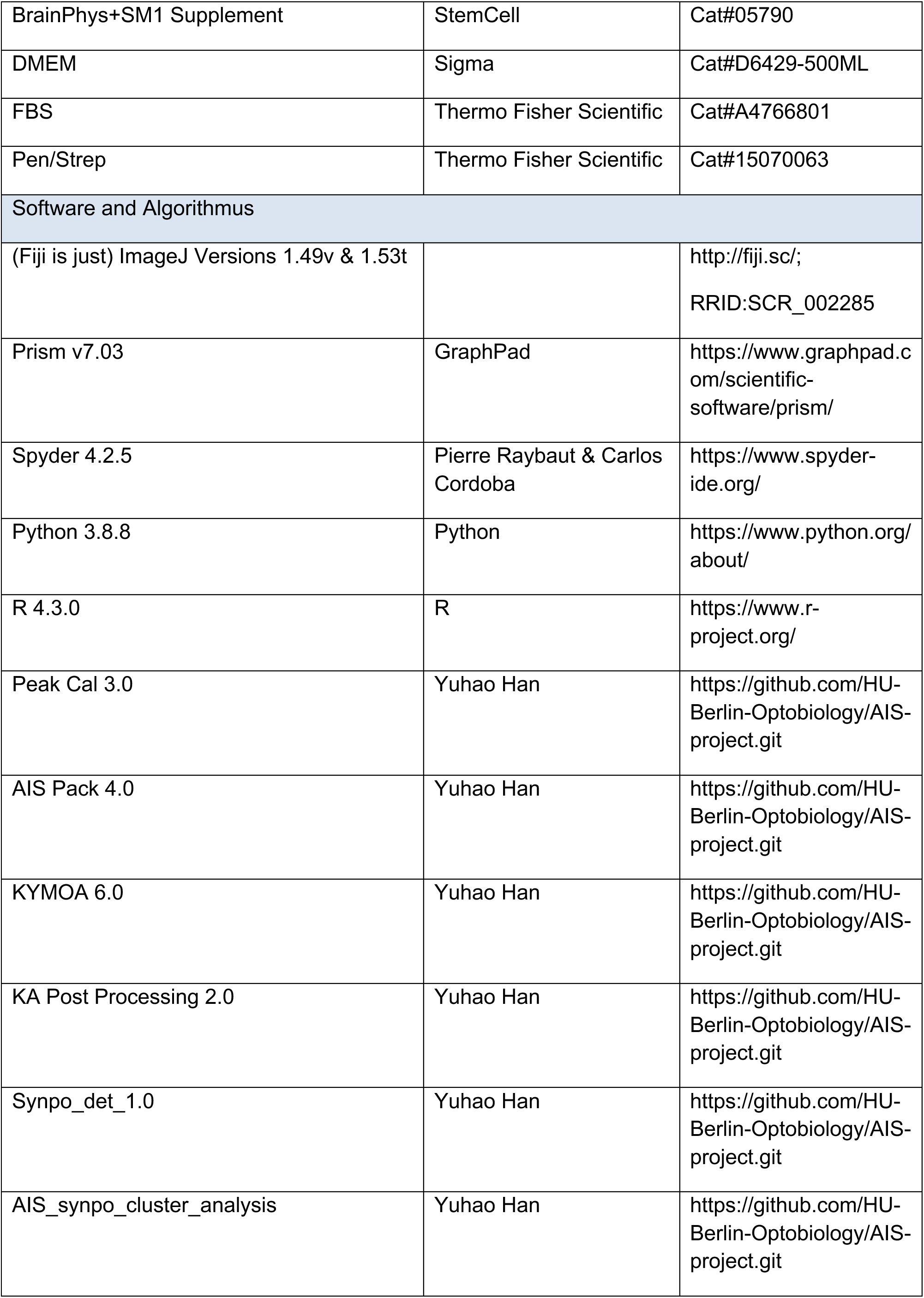

## Supplemental Videos

**Video S1.** Differentiation of nonAcD neurons. Related to Figure 1D. White arrowhead indicates the axon. Frame rate of this video is 15 frames/second.

**Video S2.** AcD neuron develop AcD via precursor neurite collateralization. Related to Figure 1D and **S1A**. Orange arrowhead indicates the precursor neurite. White arrowhead indicates the axon. Yellow and pink arrowhead indicates the AcD and stem dendrite, respectively. White arrow indicates the AIS. Black arrow indicates the cell body of AcD neuron being recorded. Frame rate of this video is 15 frames/second.

**Video S3.** AcD neuron develop axon and AcD via bifurcation of the precursor neurite. Related to **Figure S1B**. Orange arrowhead indicates the precursor neurite. White arrowhead indicates the axon. Yellow and pink arrowhead indicates the AcD and stem dendrite, respectively. White arrow indicates the AIS. Black arrow indicates the cell body of AcD neuron. Frame rate of this video is 15 frames/second.

**Video S4.** AcD neuron develop axon from collateral of the precursor neurite. Related to **Figure S1A**. Orange arrowhead indicates the precursor neurite. White arrowhead indicates the axon. Yellow and pink arrowhead indicates the AcD and stem dendrite, respectively. White arrow indicates the AIS. Black arrow indicates the cell body of AcD neuron. Frame rate of this video is 15 frames/second.

**Video S5.** MT orientation in the axon, stem dendrite and AcD of AcD neuron. Related to Figure 2A. Black arrow indicates the axon. Pink arrowhead indicates the stem dendrite. Yellow indicates the AcD. Frame rate of this video is 30 frames/second.

**Video S6.** MT orientation in the AIS and somatic dendrite of AcD neuron. Related to Figure 2A. Black arrowhead indicates the AIS. Cyan arrowhead indicates the regular somatic dendrite (Regular D). Frame rate of this video is 30 frames/second.

**Video S7.** MT orientation in nonAcD neuron. Related to Figure 2A. Black arrow indicates the axon. Black arrowhead indicates the AIS. Cyan arrowhead indicates the regular somatic dendrite (Regular D). Frame rate of this video is 30 frames/second.

**Video S8.** EGFP-Rab3A trafficking in AcD neuron. Related to Figure 4. Black arrowhead indicates the AIS. Pink arrowhead indicates the stem dendrite. Cyan arrowhead indicates the regular somatic dendrite (Regular D). Frame rate of this video is 30 frames/second.

**Video S9.** EGFP-Rab3A trafficking in nonAcD neuron. Related to Figure 4. Black arrowhead indicates the AIS. Frame rate of this video is 30 frames/second.

**Video S10.** TfR trafficking in AcD neuron. Related to Figure 5. Black arrowhead indicates the AIS. Pink arrowhead indicates the stem dendrite. Yellow arrowhead indicates the axon carrying dendrite (AcD). Black arrow indicates the axon. Frame rate of this video is 30 frames/second.

**Video S11.** TfR trafficking in nonAcD neuron. Related to Figure 5. Black arrowhead indicates the AIS. Cyan arrowhead indicates the regular somatic dendrite (Regular D). Black arrow indicates the axon. Frame rate of this video is 30 frames/second.

## Notes

### Competing Interest Statement

The authors have declared no competing interest.

### Summary of Updates

This revised version includes additional analysis of data on the abundance and size of cisternal organelles (COs) in the AIS of AcD and non-AcD neurons. Furthermore we describe the effect of chronic plasticity on the COs number. We also included new immunostainings for AIS components and found that the AIS of AcD neurons has less sodium channels.

