## Supplemental figures for "Unveiling the cell biology of hippocampal neurons with dendritic axon origin"

### Supplemental Information

#### Supplemental Figures

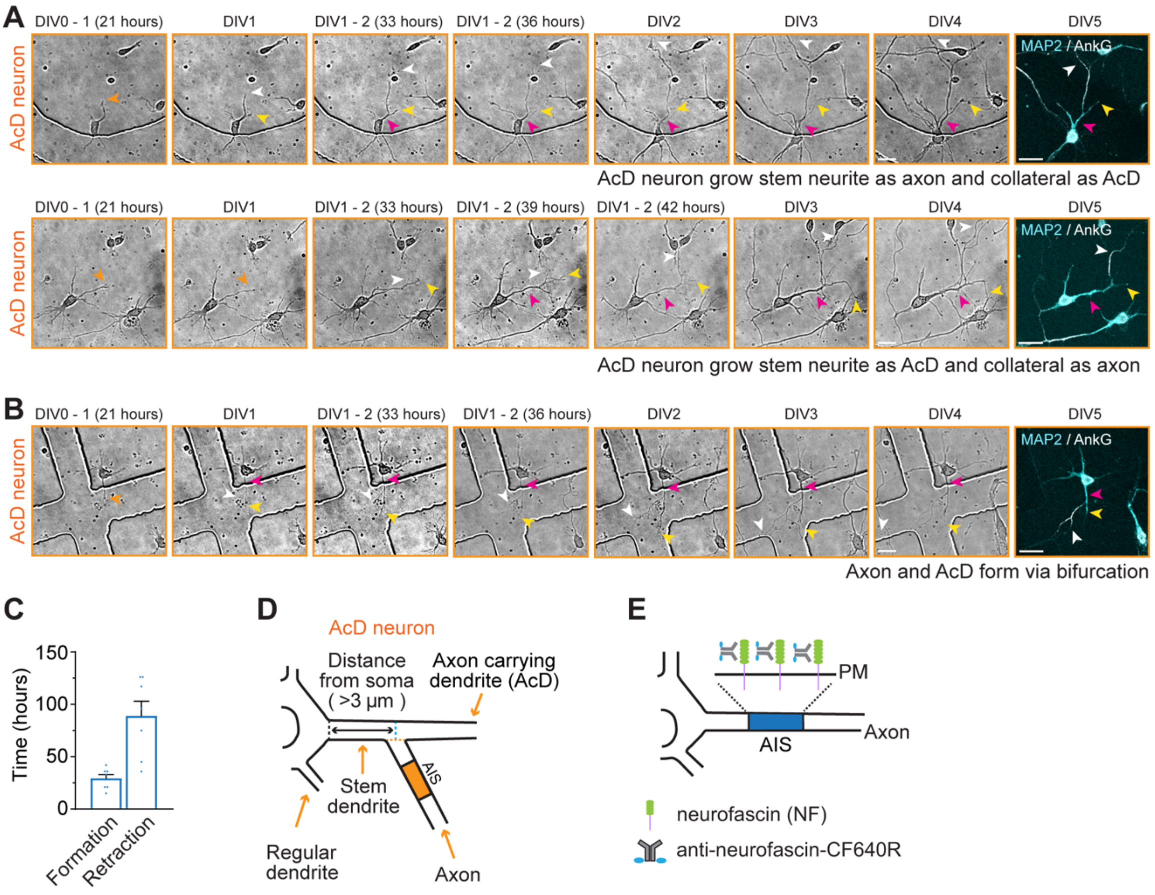


**Figure S1. Examples of atypical sequence of events during nonAcD and AcD neuron development.**

(**A**) Top row: Representative images from a time-lapse recording of AcD neuron developing a precursor neurite as axon and collateral as AcD (majority of cases). Related to **Figure 1D** and **Video S2**. Bottom row: Representative images from time-lapse recording of AcD neuron growing a precursor neurite as AcD and collateral as axon (atypical sequence). Related to **Video S4**. Orange arrowhead indicates the precursor neurite of AcD neuron. White arrowhead indicates axon. Yellow and pink arrowhead indicates AcD and stem dendrite, respectively. Dendrites and the AIS is labelled by post-staining with MAP2 and AnkG, respectively. Scale bar is 20 µm.

(**B**) Representative images from a time-lapse recording of an AcD neuron developing an axon and AcD via bifurcation of the precursor's neurite growth cone. Related to **Video S3**. Orange arrowhead indicates precursor neurite. White arrowhead indicates axon. Yellow and pink arrowhead indicates AcD and stem dendrite respectively. Cell body with dendrites and the AIS are labelled with MAP2 and AnkG antibody, respectively. Scale bar is 20 µm.

(**C**) Average time point of nonAcD neuron forming collaterals at the proximal region of the axon precursor neurite and time point at which collaterals retracted. Mean ± SEM, 1 independent culture, nonAcD: n=7 cells.

(**D**) Schematic of AcD neuron where the point of axon emergence and edge of soma are not parallel. Black dashed line indicates the edge of soma; orange dashed line indicates the border of axon start; cyan dashed line indicates the axis perpendicularly extended from the centre of axon start. Related to **Figure 1B**.

(**E**) Schematic of AIS live-labelling using a pre-conjugated antibody against extracellular domain of neurofascin (anti-NF-CF640R).

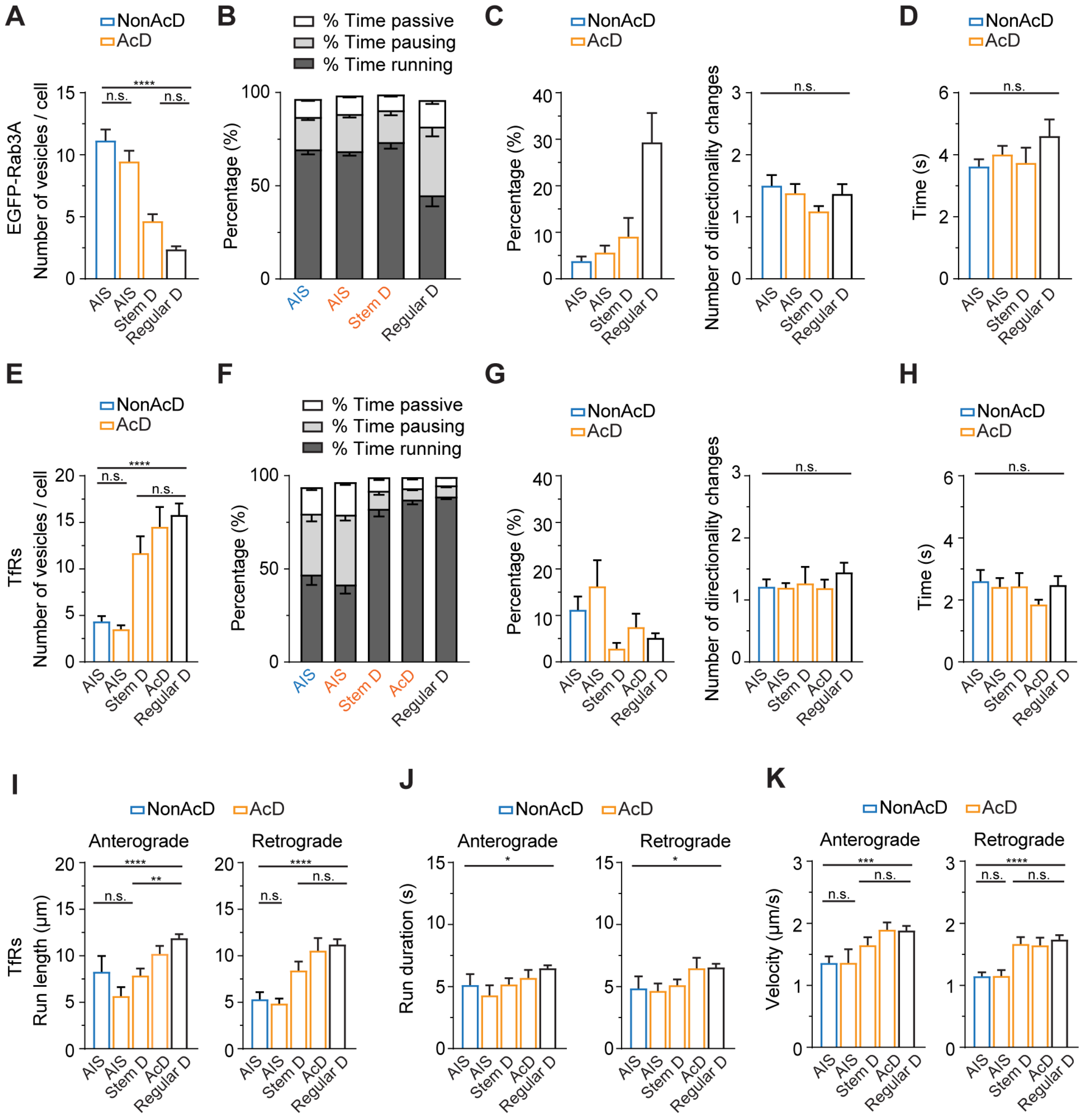


**Figure S2. Trafficking of axonal and dendritic cargoes in nonAcD and AcD neurons.**

(**A-D**) Trafficking of EGFP-Rab3A vesicles in the AIS of nonAcD neuron, the AIS and stem dendrite (Stem D) of AcD neuron, and regular dendrite (Regular D) of both nonAcD and AcD neurons. (**A**) Number of moving EGFP-Rab3 vesicles. (**B**) Percentage of time a mobile Rab3A vesicle is running, pausing or passively moving during anterograde transport. (**C**) Percentage of Rab3A vesicles change directions during anterograde transport (left) and the average number of direction changes per vesicle (right). (**D**) Average duration of each pause per EGFP-Rab3A vesicle during trafficking. Mean ± SEM, 7 independent cultures, AIS (nonAcD) n = 42 cells, AIS (AcD) n = 42 cells, Stem D n= 28 cells, Regular D n= 27 cells.

(**E-H**) Trafficking of TfR vesicles in the AIS of nonAcD neuron, the AIS, stem dendrite (Stem D) and AcD of AcD neuron, and regular dendrite (Regular D) of both nonAcD and AcD neurons. (**E**) Number of running TfR vesicles. (**F**) Percentage of time a mobile TfR vesicle is running, pausing or passively moving during anterograde transport. (**G**) Percentage of TfR vesicles change directions during anterograde transport (left graph) and the average number of direction change per vesicle (right graph). (**H**) Average duration of each pause per TfR vesicle during trafficking. Mean ± SEM, 5 independent cultures, AIS (nonAcD) n = 25 cells, AIS (AcD) n = 23 cells, Stem D n= 23 cells, AcD n = 19 cells, Regular D n= 56 cells.

(**I**, **J** and **K**) Average length (**I**), duration (**J**) and velocity (**K**) of TfR vesicles running towards anterograde and retrograde directions in the AIS of nonAcD neuron, the AIS, stem dendrite (Stem D) and AcD of AcD neuron, and the regular dendrite (Regular D) of both nonAcD and AcD neuron. Mean ± SEM, 5 independent cultures, Anterograde: AIS (nonAcD): n = 21 cells, AIS (AcD) n = 15 cells, Stem D: n= 23 cells, AcD: n = 19 cells, Regular D: n= 54 cells, Retrograde: AIS (nonAcD) n = 23 cells, AIS (AcD) n = 23 cells, Stem D n= 20 cells, AcD n = 18 cells, Regular D n= 56 cells.

One-way ANOVA with Tukey's multiple comparisons test, no significance (n.s.) $p > 0.05$, ****$p < 0.0001$.

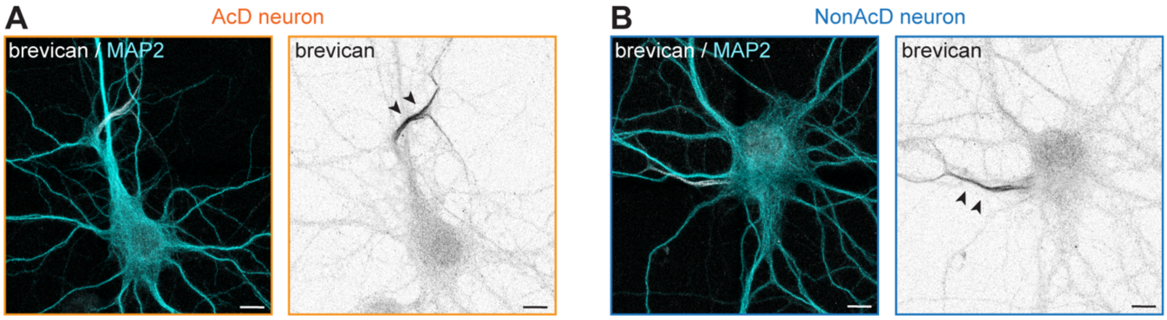


**Figure S3. Similarly to nonAcD neurons, AcD neurons form specific ECM at the AIS.**

(**A** and **B**) Representative images of AIS-specific ECMs labelled by brevican staining at the AIS of DIV21 nonAcD and AcD neurons. Black arrowheads indicate brevican at the AIS. Scale bar is 10 μm.

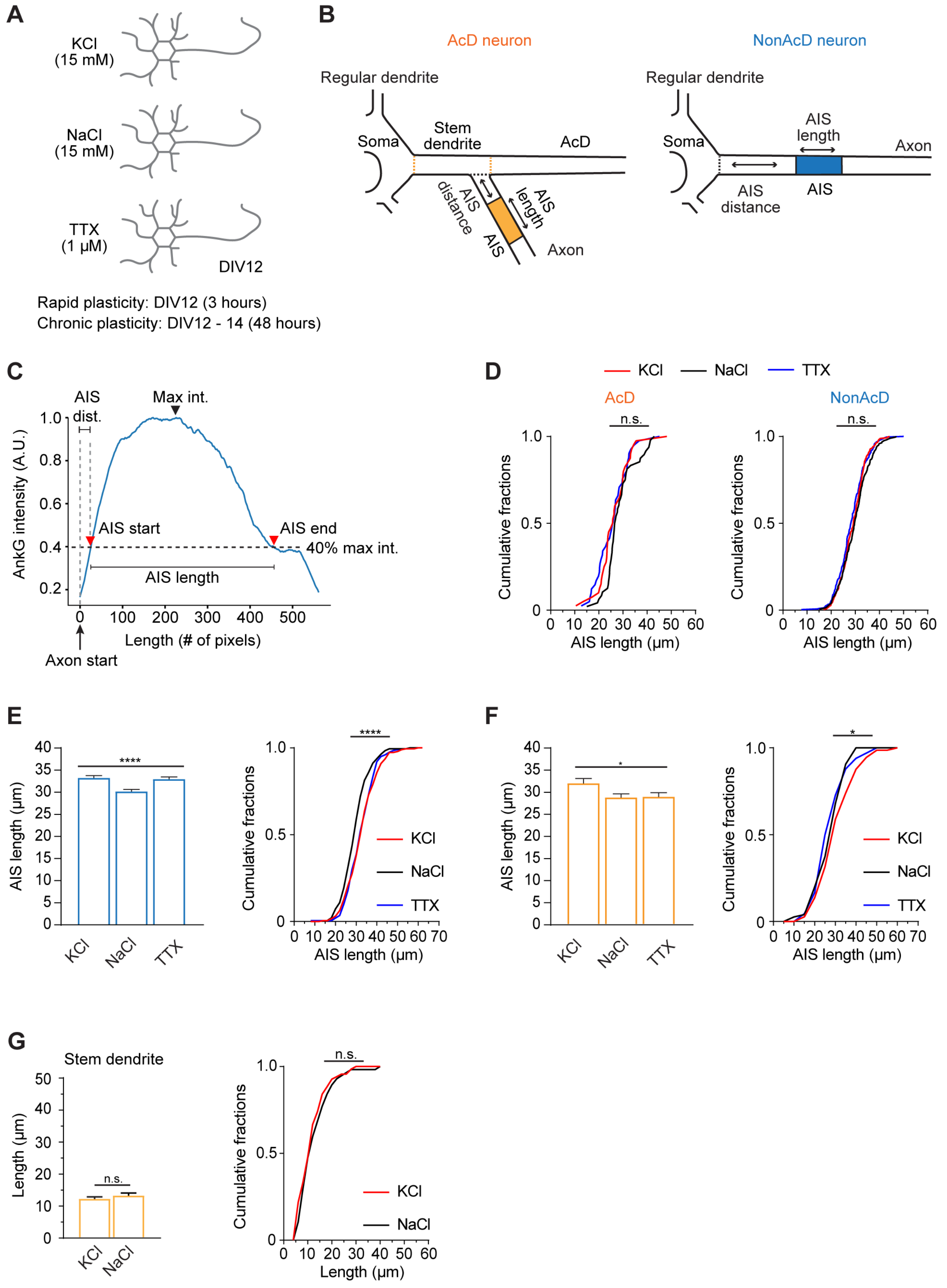


**Figure S4. Definition of AIS length and distance from the cell body or axon carrying dendrite and illustration of the workflow to induce AIS plasticity.**

(**A**) Schematic of AIS rapid and chronic plasticity experiment.

(**B**) Definition of AIS length and AIS distance in nonAcD and AcD neurons as read out of AIS plasticity. Orange dashed line indicates the border of stem dendrite of AcD neurons; black dashed line indicates the border of axon origin.

(**C**) Illustration of intensity-based AIS length and AIS distance measurement. AIS dist. stands for AIS distance.

(**D**) Cumulative of AIS length in nonAcD and AcD neuron upon induction of rapid plasticity. Mean ± SEM, 3 independent cultures, nonAcD: n (KCl) = 298 cells, n (NaCl) = 264 cells, n (TTX) = 306 cells, AcD: n (KCl) = 48 cells, n (NaCl) = 40 cells, n (TTX) = 41 cells.

(**E** and **F**) Measurement of AIS length in nonAcD (**E**) and AcD (**F**) neurons upon induction of chronic plasticity. Mean ± SEM, 4 independent cultures, nonAcD: n (KCl) = 189 cells, n (NaCl) = 192 cells, n (TTX) = 202 cells, AcD: n (KCl) = 73 cells, n (NaCl) = 75 cells, n (TTX) = 66 cells.

(**G**) Measurement of stem dendrite length in AcD neuron upon induction of chronic plasticity. Mean ± SEM, 4 independent cultures, AcD: n (KCl) = 73 cells, n (NaCl) = 75 cells, n (TTX) = 66 cells. Mann-Whitney test, no significance (n.s.) $p > 0.05$.

One-way ANOVA with Tukey's multiple comparisons test, no significance (n.s.) $p > 0.05$, *$p < 0.05$, ****$p < 0.0001$.

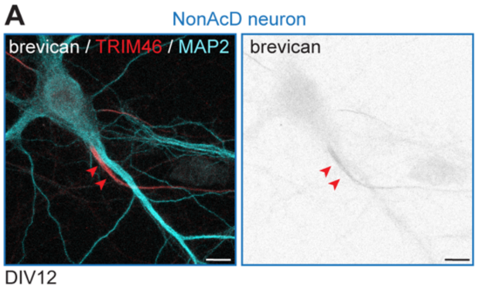


**Figure S5. Expression of AIS specific ECM protein brevican in DIV12 neurons**

(**A**) Immunostaining of DIV12 neurons with AIS specific ECM protein brevican. Red arrowheads indicate the AIS. Scale bar is 10 μm.

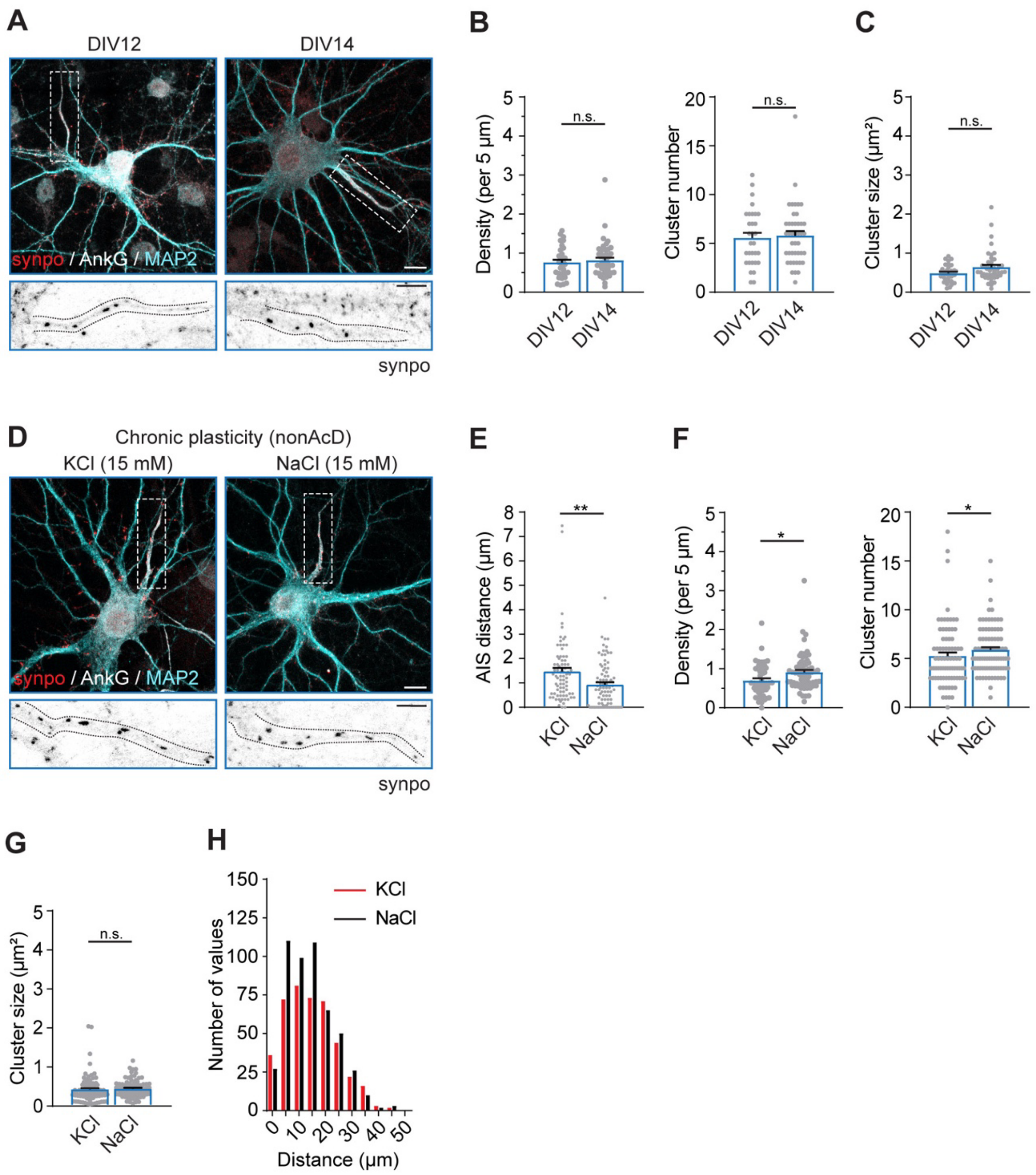


**Figure S6. Increased neuronal activity reduces the number of COs at the AIS.**

(**A**) Representative images of DIV12 and DIV14 neurons with labelled CO (synpo), the AIS (AnkG) and the somato-dendritic compartment (MAP2). Scale bar is 10 µm. Inset: Zoom-ins corresponding to white dashed rectangular. Black dashed lines indicate the edge of AIS. Scale bar is 10 µm.

(**B**) Quantification of synpo cluster density (left) and number (right) at the AIS of DIV12 and DIV14 neurons. Mean ± SEM, 3 independent cultures, DIV12: n= 31 cells, DIV14: n= 43 cells.

(**C**) Quantification of synpo cluster size at the AIS of DIV12 and DIV14 neurons. Mean ± SEM, 3 independent cultures, DIV12: n= 31 cells, DIV14: n= 43 cells.

(**D**) Representative images of DIV14 neurons with labelled CO (synpo), the AIS (AnkG) and the somato-dendritic compartment (MAP2). Neurons were treated with KCl and NaCl from DIV12 for 48 hours to induce chronic plasticity. Scale bar is 10 µm. Inset: Zoom-ins corresponding to white dashed rectangular. Black dashed lines indicate the edge of AIS. Scale bar is 10 µm.

(**E**) AIS distance from the soma in nonAcD neurons upon induction of chronic plasticity. Mean ± SEM, 3 independent cultures, n (KCl) = 79 cells, n (NaCl) = 86 cells.

(**F**) Quantification of synpo cluster density (left) and number (right) at the AIS of DIV14 neurons upon induction of chronic AIS plasticity. Mean ± SEM, 3 independent cultures, KCl: n= 80 cells, NaCl: n= 86 cells.

(**G**) Quantification of synpo cluster size at the AIS of DIV14 neurons upon induction of chronic AIS plasticity. Mean ± SEM, 3 independent cultures, KCl: n= 77 cells, NaCl: n= 87 cells.

(**H**) Distribution of synpo clusters at the AIS of nonAcD neurons upon induction of chronic plasticity. In total 3 independent cultures, n (KCl) = 420 clusters from 79 cells, n (NaCl) = 501 clusters from 86 cells.

Mann-Whitney test: not significant (n.s.) p > 0.05, *p < 0.05, **p < 0.001, ***p < 0.0001.
